# Predictable adaptive evolution of a phage endolysin through substrate recognition optimization

**DOI:** 10.64898/2026.05.06.723369

**Authors:** Xiaojun Zhu, Vincent Somerville, Rong Shi, Rojo V. Rakotoharisoa, Roberto A. Chica, Sylvain Moineau, Frank Oechslin

## Abstract

Bacteriophages release progeny by producing endolysins that degrade the bacterial cell wall. While the frequent horizontal transfer of endolysins suggests substantial evolutionary plasticity, the mechanisms by which these enzymes adapt to new phage-host contexts remain poorly understood. Here, we investigated the evolutionary dynamics and structural mechanisms governing endolysin adaptation following experimental evolution of a chimeric phage generated via heterologous endolysin exchange between phages infecting different hosts.

Using replicate experimental evolution and time-resolved PacBio sequencing, we identified a dominant, highly reproducible adaptive trajectory characterized by the stepwise fixation of three key mutations. This constrained mutational order correlated with incremental gains in enzymatic activity, reflecting a rugged yet predictable fitness landscape. High-resolution structural analyses revealed that these substitutions lie exclusively outside the catalytic site; instead, they enhance substrate recognition through electrostatic tuning, optimized hydrophobic packing, and local conformational refinement, resulting in significantly higher binding affinity.

While the adaptive trajectory was largely conserved, one replicate followed an alternative path, highlighting the interplay between selection and historical contingency. Adaptation was further shaped by a functional trade-off, whereby increased lytic activity on the novel host was accompanied by reduced activity on the ancestral host, consistent with antagonistic pleiotropy. Genome-wide sequencing additionally identified a compensatory mutation in a lytic transglycosylase, suggesting coordinated evolution of the broader lysis machinery. Together, these results demonstrate that endolysins evolve through reproducible adaptive walks constrained by structure, selection, and trade-offs, providing a mechanistic framework for understanding enzyme evolution and informing rational protein engineering.

## Introduction

Endolysins are enzymes produced by (bacterio)phages to breach the bacterial cell wall at the end of their replication cycle, facilitating the release of progeny virions ^1^. Their potent bacteriolytic activity has drawn considerable attention as a basis for antibacterial therapy, particularly against multidrug-resistant Gram-positive pathogens ^2^. These enzymes specifically target peptidoglycan, the primary structural component of the bacterial cell wall, composed of glycan chains cross-linked by short stem peptides ^3^. Endolysins contain enzymatically active domains (EADs) that are classified into four main groups based on their target bonds: glucosaminidases and muramidases, which hydrolyze glycan chains, and amidases and endopeptidases, which act on stem peptides or cross-linking bridges ^4^. In Gram-positive systems, endolysins often feature a modular architecture, with one or more cell wall-binding domains (CBDs) linked to the EAD via short connectors ^4^. These non-catalytic domains facilitate adsorption to specific cell wall components, such as teichoic acids or peptidoglycan units, bringing the catalytic domain into proximity with its substrate and enhancing lytic efficiency ^5–7^.

The modular recombination of EADs and CBDs, fueled by frequent horizontal gene transfer, has generated hundreds of structurally distinct endolysins ^8–11^. This diversity reflects intense selective pressures as phages adapt to the structural nuances of different bacterial cell walls ^12^. Because peptidoglycan variation is significantly greater between species than within species, diversifying selection acts most strongly on the endolysins of phages infecting phylogenetically distant hosts ^13^. Supporting this, experimental swapping of structurally distinct endolysin genes between phages has been shown to impose substantial fitness costs when phages encounter distantly related strains or species ^14^.

Although studies generating chimeric endolysins in vitro have demonstrated that domain shuffling can alter host range ^15–17^, the micro-evolutionary processes by which these enzymes refine their specificity in natural contexts remain poorly understood. We have previously shown that endolysins can rapidly acquire adaptive mutations when introduced into a phage infecting a novel host ^14^, and can further adapt to environmental conditions ^18^, underscoring their remarkable evolutionary plasticity. Despite these observations, the molecular mechanisms by which EADs and CBDs accommodate diverse cell wall structures while maintaining catalytic activity—and how adaptive mutations remodel the enzyme–substrate interface—remain largely unexplored.

To address these adaptive processes in detail, we leveraged a chimeric phage previously generated via heterologous endolysin exchange between phages infecting different host strains ^14^. By tracking mutations over multiple independent evolutionary trajectories and linking these genomic changes to high-resolution structural and enzymatic data, we uncover principles governing how adaptive changes in non-catalytic regions modulate substrate engagement and specificity, balancing activity between novel and ancestral hosts. These findings provide general insights into the evolution of enzyme function and inform the rational design of phage-derived proteins for therapeutic and biotechnological applications.

## Results

### In vivo evolution of a phage endolysin toward a novel host

Building on previous work showing that recombinant phages carrying heterologous endolysins can rapidly adapt to novel hosts ^14^, we used this system to dissect the molecular and structural basis of endolysin adaptation. We employed a chimeric derivative of the lactococcal phage P008, P008::Lys1358, in which the native P008 endolysin (LysP008) was replaced by Lys1358, the endolysin of the distantly related, narrow-host-range lactococcal phage 1358 ^14,19^ (Fig. 1a). The two phages infect non-overlapping *Lactococcus lactis* strains: P008 targets IL1403 ^20^, whereas phage 1358 infects strain 582 ^21^, which reaches maximal growth later (454.3 ± 5.8 vs. 307.7 ± 5.8 min; Supplementary Fig. 1) and has a thicker cell wall (19.2 ± 1.5 vs. 13.6 ± 1.3 nm; P < 0.001; Fig. 1b, Supplementary Fig. 2). Structurally, LysP008 contains an N-terminal amidase, which cleaves the bond between N-acetylmuramic acid and L-alanine of the stem peptide ^22^, and a C-terminal LysIME-EF1L-like domain ^23^. In contrast, Lys1358 has an N-terminal CHAP domain, reported to cleave peptide bonds in peptidoglycan stems or cross-bridges ^24^, and a C-terminal SH3b-type CBD that recognizes specific peptidoglycan motifs or polysaccharides ^6,25,26^ (Fig. 1a).

**Fig. 1.**
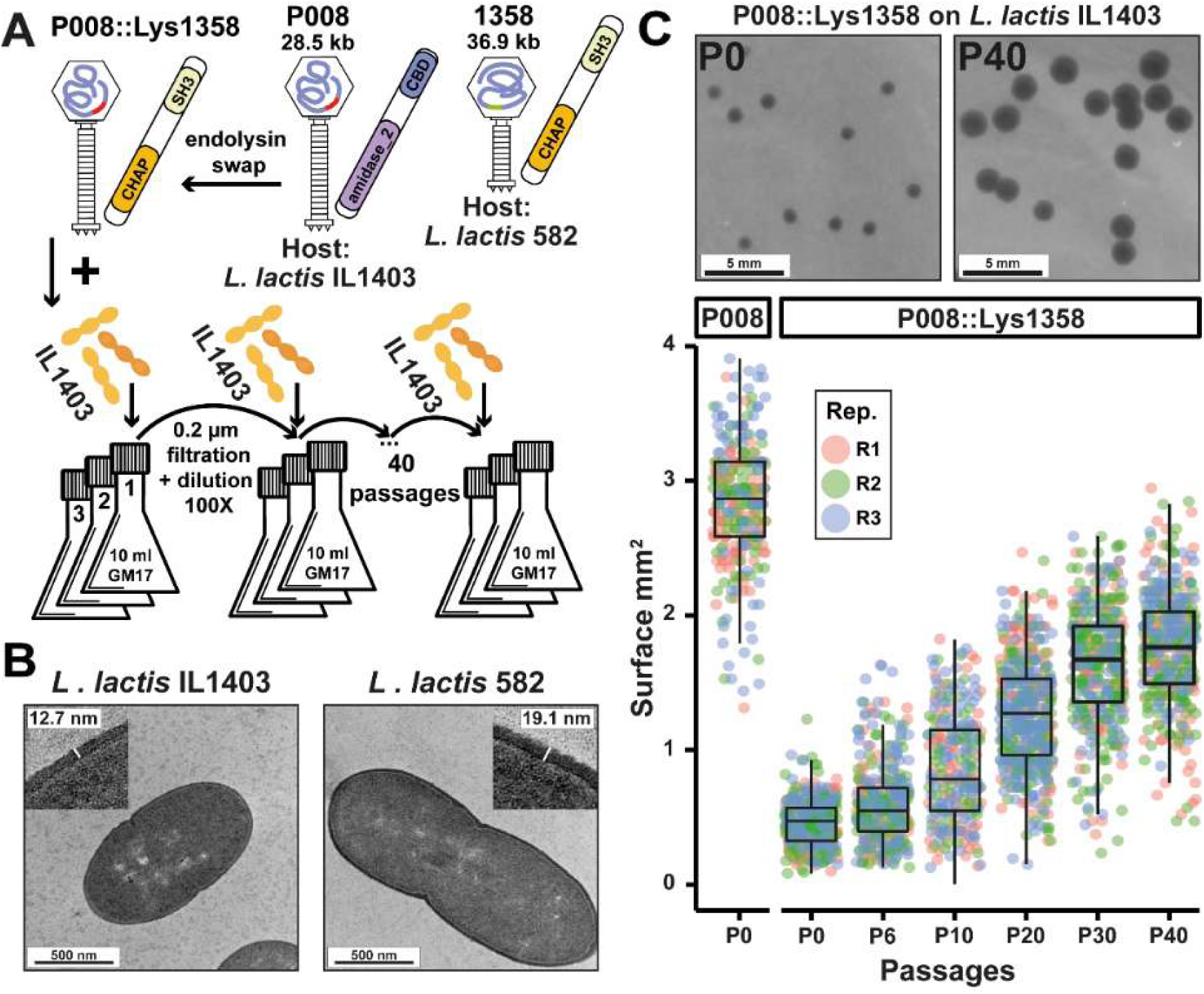
Serial adaptation of a recombinant lactococcal phage carrying a heterologous endolysin. **(a)** Experimental evolution of the chimeric phage. Schematic of the recombinant phage P008::Lys1358, constructed by replacing the native LysP008 with Lys1358 from the distantly related phage 1358, as established previously 14. This chimeric phage was used as the starting point for 40 serial passages on *L. lactis* IL1403 in triplicate to drive endolysin adaptation. **(b)** Host cell wall ultrastructure. Representative transmission electron microscopy (TEM) images of *L. lactis* strains IL1403 and 582, highlighting the significantly thicker cell wall of strain 582 compared to IL1403 (19.2 ± 1.5 vs. 13.6 ± 1.3 nm; P < 0.001). **(c)** Phenotypic gain in lytic fitness. Representative plaque morphologies illustrate the expansion of clear zones from the onset (passage 0) to the conclusion (passage 40) of the experiment. Consistent with these visual changes, quantification of plaque areas at indicated intervals (passages 6–40) reveals a progressive increase in lytic activity, ultimately surpassing both the initial recombinant and wild-type P008.

To drive adaptation, P008::Lys1358 was serially passaged over 40 passages on IL1403 cells in triplicate (Fig. 1a). At each passage, host cells were replaced with fresh IL1403, allowing only the recombinant phage to evolve. At passage 0 (P0), P008::Lys1358 formed significantly smaller plaques than wild-type P008 (0.46 ± 0.17 vs. 2.83 ± 0.46 mm²; Wilcoxon rank-sum test, p < 0.0001; Figs. 1b, c). Plaque size increased steadily over the experiment (0.035 mm² per passage; linear regression, p < 0.0001), reflecting progressive gains in lytic activity on the new host.

### Convergent molecular trajectories during endolysin host adaptation

To resolve the positions, frequencies, and linkage (synteny) of mutations accumulating during Lys1358 adaptation, we performed full-length deep sequencing of the *lys1358* gene across all replicates at two-passage intervals using PacBio long-read technology. Based on the baseline error rate and genetic diversity of the isogenic starting population (P₀), we set a 1% frequency threshold for reliable de novo mutation detection.

We identified three recurrent nonsynonymous mutations across the CHAP and SH3b domains in all three independent lineages, suggesting strong convergent selection (Fig. 2a). The first substitution, N66D (Asn66Asp; 196A>G) within the CHAP domain, emerged by passage 6 and reached fixation by passage 20 across all replicates (Fig. 2b). This was followed by the emergence of S167I (Ser167Ile; 500G>T) in the SH3b domain, which reached fixation at replicate-dependent rates. A third SH3b mutation, L186I (Leu186Ile; 556C>A), emerged later in the experiment but remained polymorphic, never reaching fixation within the 40-passage window (Fig. 2b).

**Fig. 2.**
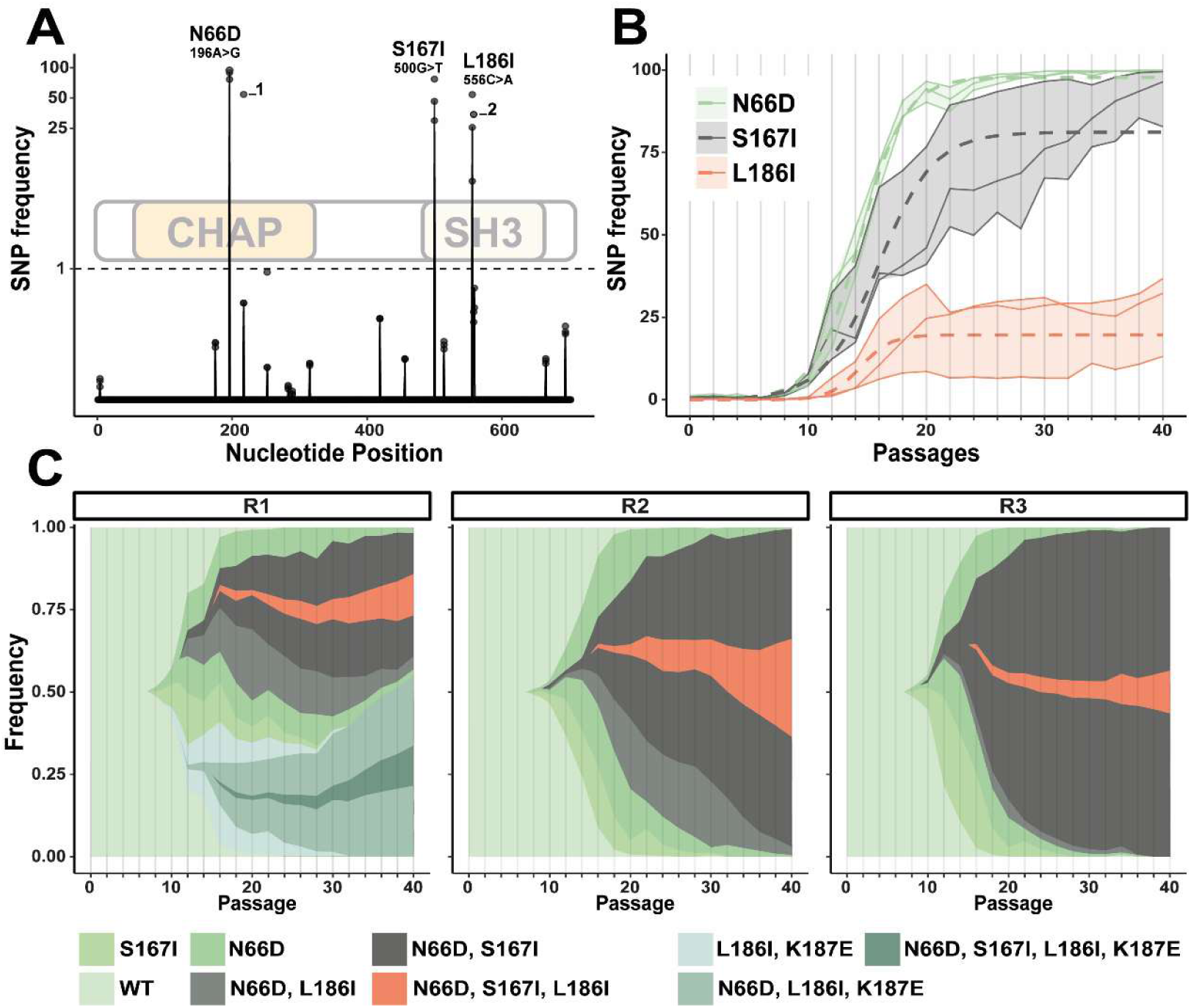
Evolutionary trajectories and successional dynamics of *lys1358* mutations. **(a)** Mutational landscape of *Lys1358*. Frequency of all single-nucleotide polymorphisms (SNPs) identified across the three independent replicates (points) and their mean values (solid line) at passage 40. The *de novo* mutation detection threshold (1%) is indicated by a dashed horizontal line. The structural architecture of the endolysin, comprising the N-terminal CHAP and C-terminal SH3b domains, is indicated. The three primary adaptive mutations N66D, S167I, and L186I are labeled. The synonymous mutation Q73Q (217G>A) and the non-synonymous mutation K187E (559A>G), which appeared in only one replicate, are labeled as 1 and 2, respectively. **(b)** Frequency trajectories of focal SNPs. Dynamics of the three adaptive mutations N66D, S167I, and L186I over 40 passages. Individual replicates are represented by solid lines with corresponding shaded areas. The average fit for each SNP is shown as a dashed line (Gompertz growth model). Vertical lines indicate PacBio sequencing timepoints. **(c)** Haplotype dynamics. Frequencies of specific mutational combinations (linkage groups) across the three replicates. This panel illustrates the successional displacement of single variants by double and triple mutants over the course of the experiment.

The temporal appearance of these mutations was reflected in their distinct frequency trajectories and successional linkage patterns (Fig. 2c). While N66D single variants were widespread early in the experiment, S167I was rarely detected as a single variant, and L186I single variants were never observed. Instead, the N66D single-variant populations were rapidly displaced by double-variant selective sweeps, predominantly the N66D + S167I genotype. In the later stages of evolution, triple variants (N66D + S167I + L186I) emerged and expanded, indicating an additive fitness benefit in which subsequent mutations were favored only on the adapted N66D background.

Lineage-specific variations provided further insight into the mutational landscape. In replicate 1, a nonsynonymous mutation, K187E (Lys187Glu; 559A>G), consistently co-occurred with L186I, following the broader successional pattern of association with the N66D and S167I backgrounds. Additionally, a synonymous substitution at Q73Q (Gln73Gln, 217G>A) was detected in replicate 2; its stable frequency alongside N66D suggests it reached high frequency through genetic hitchhiking rather than direct selection (Fig. 2a).

### Adaptive substitutions enhance lytic activity and substrate binding while inducing host-specific trade-offs

To assess the functional impact of the adaptive substitutions N66D, S167I, and L186I, we purified wild-type Lys1358 and variants carrying individual or combined substitutions (Supplementary Fig. 3). We first quantified lytic activity against *L. lactis* IL1403, the host used during the experimental evolution of phage P008::Lys1358 (Fig. 3a). Activity was analyzed using a linear mixed-effects model with variant and concentration as fixed effects and biological replicate as a random effect. Pairwise comparisons derived from estimated marginal means (EMMs; Supplementary Table 1) revealed concentration-dependent effects on lytic activity.

**Figure 3.**
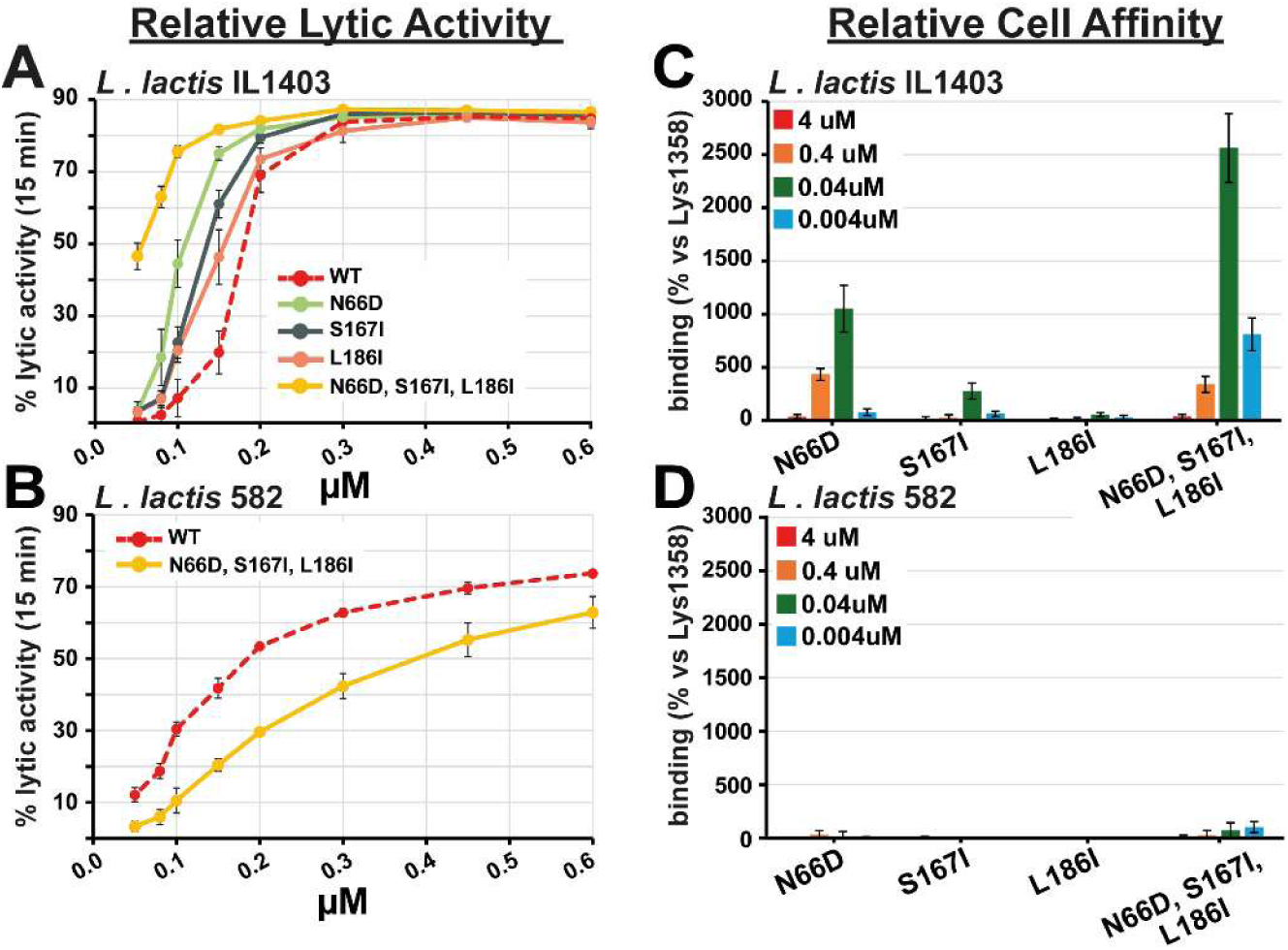
Adaptive mutations in Lys1358 enhance lytic efficiency and binding affinity, driving host specificity. **(a)** Dose-dependent lytic activity on the selection host (*L. lactis* IL1403). Lytic activity of Lys1358 and its derived variants (N66D, S167I, L186I, and the triple variant) against *L. lactis* IL1403, the strain used for experimental evolution of phage P008::Lys1358. Activity is expressed as the percentage reduction in turbidity (OD₆₀₀) after 15 min relative to T₀. Results are shown across a range of enzyme concentrations (0.05–0.6 µM). **(b)** Evaluation of host-switch trade-offs. Comparison of lytic activity between WT Lys1358 and the triple variant on the ancestral host (*L. lactis* 582), the original host of phage 1358 from which Lys1358 was identified. **(c, d)** Comparative cell-binding affinity. Relative cell binding affinity of catalytically inactive variants (C29A) to surface-adsorbed *L. lactis* IL1403 (c) and *L. lactis* 582 (d) was quantified by ELISA. Cell surface bound enzymes were detected using anti-6×His primary and HRP-conjugated secondary antibodies. Affinity values are normalized to WT Lys1358 (100%) at each concentration to facilitate comparison of mutational effects. All experimental measurements **(a–d)** were performed in ≥3 replicates using two independently purified protein batches.

At 0.05 µM, only the triple variant exhibited significantly higher activity than Lys1358 (*P* < 0.01). In the intermediate concentration range (0.08–0.2 µM), all variants showed increased activity relative to Lys1358, although the magnitude and statistical support varied with concentration. N66D, S167I, and the triple variant showed strong and consistent increases (typically *P* < 0.0001), whereas L186I exhibited more modest effects. The triple variant consistently showed the largest effect, while N66D conferred the strongest increase among single substitutions, reaching approximately fivefold higher activity at 0.1 µM. At higher concentrations (0.3–0.6 µM), no significant differences were observed (*P* > 0.05), consistent with enzyme saturation. Among single variants, activity differences followed the order N66D > S167I > L186I, mirroring their sequential emergence during experimental evolution (Supplementary Table 2). This hierarchy was further supported by pairwise comparisons (Supplementary Table 3), which confirmed significant differences at 0.15 µM (L186I vs. N66D: estimate = –28.72, *P* < 0.0001; L186I vs. S167I: estimate = –14.65, *P* < 0.0001).

To assess potential evolutionary trade-offs, the lytic activities of Lys1358 and the triple variant were compared on the ancestral host (*L. lactis* 582). The triple variant showed significantly reduced activity on strain 582 compared to Lys1358 across all tested concentrations (0.05–0.6 µM; *P* < 0.0001; Supplementary Table 4). These findings indicate a clear trade-off, whereby adaptation to the IL1403 host environment compromises enzymatic performance on the ancestral host.

We further hypothesized that increased lytic activity resulted from altered substrate recognition. To test this, catalytically inactive variants (carrying the CHAP domain mutation C29A) were generated to decouple binding from cell lysis (Supplementary Fig. 4). ELISA-based assays revealed enhanced binding affinity of the variants to *L. lactis* IL1403 cells (Fig. 3c,d; Supplementary Table 5). Variations in binding affinity were analyzed using linear mixed-effects models with pairwise comparisons of estimated marginal means (EMMs) relative to Lys1358. At 0.4 µM, binding approached saturation, whereas differences were strongly amplified at 0.004 µM. The triple variant showed the largest increase in affinity (2562% relative to Lys1358; P < 0.001), followed by N66D (1052%; P < 0.001) and S167I (278%; P < 0.001), while L186I exhibited a positive trend toward increased affinity (55%; P > 0.05), though this effect did not reach statistical significance at the concentrations tested. This clear ranking in relative binding affinities reflects the successional emergence of the mutations and directly correlates with their observed effects on lytic activity. In contrast, affinity for the ancestral host (strain 582) remained largely unchanged (Supplementary Table 6), supporting the conclusion of host-specific adaptation.

### The N66D mutation remodels the electrostatic landscape of Lys1358 without altering its overall structure

To investigate how the N66D substitution facilitates host adaptation, we determined the crystal structures of wild-type (WT) Lys1358 and the N66D variant at 1.33 Å and 1.52 Å resolution, respectively (PDB IDs: 10VD, 10VE; Supplementary Fig. 5, Supplementary Table 7). The enzyme adopts a canonical CHAP domain fold with a catalytic core highly conserved relative to the CHAP domains of the autolysin PcsB and the phage endolysin CHAPk (Cα RMSDs of 2.83 Å and 0.77 Å, respectively) ^27,28^, and contains an active site defined by the catalytic triad Cys29, His89, and Glu105 (Fig. 4a, Supplementary Fig. 6).

**Fig. 4.**
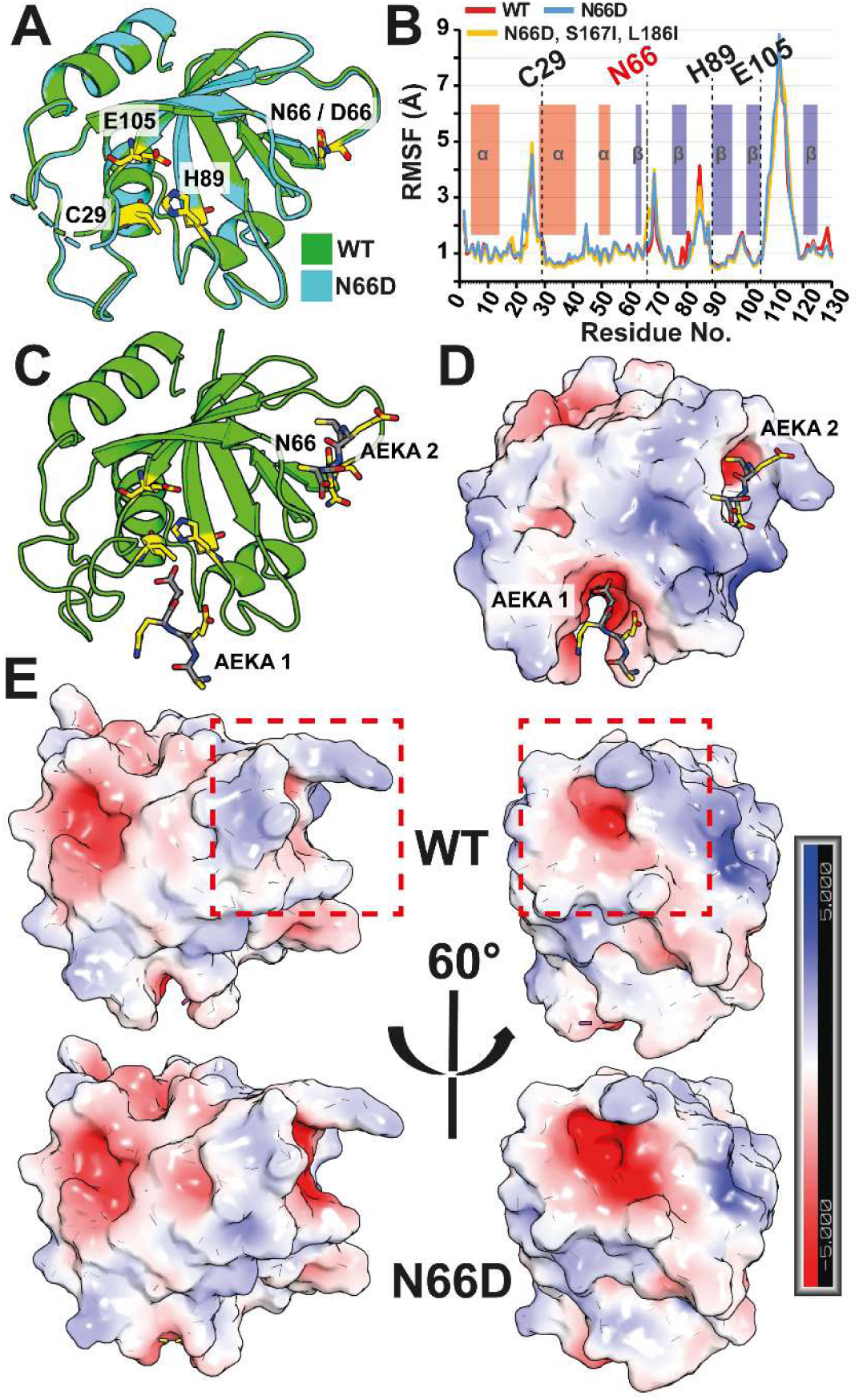
Impact of the N66D mutation on the structure and electrostatics of the Lys1358 CHAP domain. **(a)** Structural alignment of the WT Lys1358 CHAP domain (blue; 1.33 Å, PDB ID: 10VD) and the N66D variant (cyan; 1.52 Å, PDB ID: 10VE), showing high overall similarity (*Cα* RMSD = 0.13 Å). Residues 108–115 are not resolved in the crystal structures. **(b)** Root-mean-square fluctuation (RMSF) profiles derived from molecular dynamics simulations of WT Lys1358 (red), the N66D variant (blue), and the triple variant N66D/S167I/L186I (yellow). Analysis is shown for residues 1–130, with secondary structure elements (α-helices in red; β-strands in blue) indicated above. **(c)** Predicted structural complex of the CHAP domain with an AEKA peptide corresponding to the *L. lactis* peptidoglycan stem, generated using AlphaFold3. The model indicates that the peptide is accommodated within a peripheral cleft encompassing the N66 site. The peptide is shown as a grey backbone with side chains colored by atom type; catalytic residues C29 and H89, as well as the mutation site N66, are highlighted. **(d)** Electrostatic surface representation of the predicted complex shown in panel (c). Red and blue indicate negatively and positively charged regions, respectively (scale from –5 to +5 kBT/e). **(e)** Comparative electrostatic surface potential of the WT Lys1358 CHAP domain and the N66D variant. The boxed region highlights the localized enhancement of negative electrostatic potential within the peripheral cleft induced by the N66D substitution.

Notably, the N66D substitution is located on the protein surface and is physically distal from the active site. Structural alignment of the WT and N66D CHAP domains revealed no global conformational changes (Cα RMSD = 0.13 Å over 112 residues), indicating that the substitution does not perturb the overall fold or the positioning of the catalytic residues (Fig. 4a). These observations suggest that its functional impact could be mediated by altered surface properties, such as electrostatic remodeling, rather than structural reorganization.

Molecular dynamics (MD) simulations corroborated the global structural stability of the N66D variant, while root-mean-square fluctuation (RMSF) analysis identified four discrete regions of increased conformational plasticity (Fig. 4b). Notably, three of these regions (residues 24–27, 83–85, and 108–117) flank the catalytic triad, suggesting a role in optimizing active-site dynamics. The fourth region encompasses the N66 site itself, implicating localized backbone mobility at this position in facilitating substrate engagement.

To explore potential substrate interactions, we used AlphaFold3 to model the complex between the CHAP domain and the *L. lactis* peptidoglycan stem peptide (L-Ala–D-Glu–L-Lys–D-Ala; AEKA) ^13^. The predicted models suggest two distinct binding modes: one peptide occupies the primary catalytic groove, whereas a second can be accommodated within a peripheral cleft that includes N66 (Fig. 4c, Supplementary Fig. 7). Surface electrostatic potential mapping further highlights the impact of the N66D substitution. Positioned within this cleft, replacement of the neutral asparagine with a negatively charged aspartic acid increases the local negative potential (Fig. 4d, e). This change is accompanied by a decrease in the predicted isoelectric point (pI) of the CHAP domain (from ∼8.8 to ∼7.9), reflecting an overall shift in net charge. Together, these observations indicate that the mutation reshapes the electrostatic landscape of this region and may influence interactions with peptidoglycan substrates.

Finally, differential scanning fluorimetry showed that the N66D mutation is destabilizing, lowering the melting temperature from 55.5 °C to 50.5 °C (Supplementary Fig. 8). This decrease in stability is consistent with an evolutionary trade-off, in which enhanced host-specific activity may be accompanied by reduced protein robustness ^29^.

### Adaptive mutations in the SH3b domain map to key substrate recognition sites

To investigate how the S167I and L186I mutations affect peptidoglycan binding, we compared the SH3b domain of Lys1358 with that of lysostaphin, a well-characterized glycyl-glycine endopeptidase from *Staphylococcus simulans* ^30^. Lysostaphin recognizes both the stem peptide and the pentaglycine cross-bridge of *Staphylococcus aureus* peptidoglycan and has been structurally characterized in complex with synthetic analogs (PDB ID: 6RK4) ^25^. The SH3b domains of both enzymes are highly homologous (Cα RMSD of 0.76 Å over 54 aligned residues; Fig. 5a), with the primary structural divergence being a β-hairpin (residues 478–485 in lysostaphin) that is truncated in Lys1358.

**Fig. 5.**
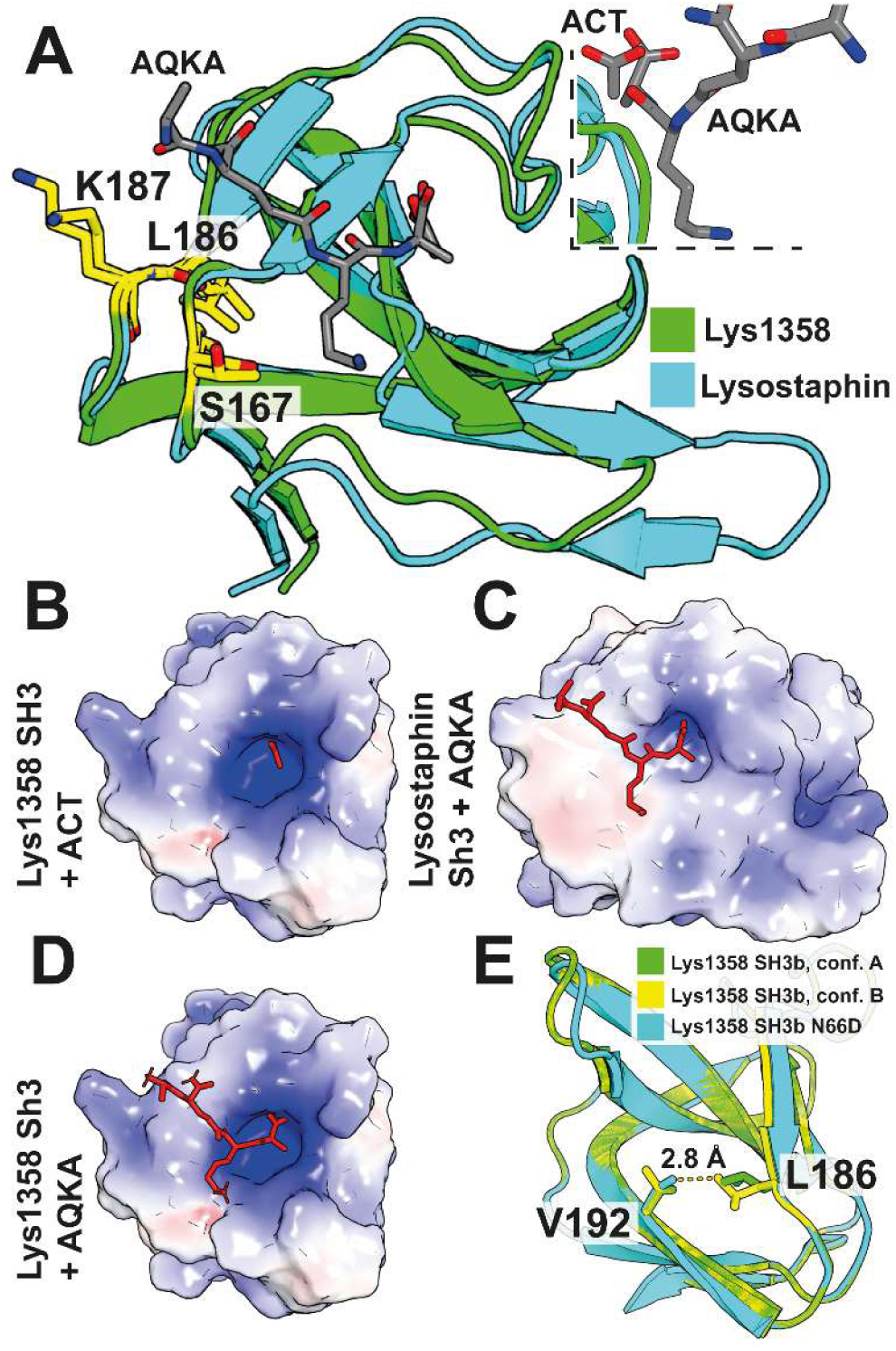
Lys1358 SH3b domain adopts a conserved fold for stem peptide recognition. **(a)** Structural superposition of the Lys1358 SH3b domain (green; residues 159–233) and the lysostaphin SH3b domain (cyan; residues 415–493; PDB ID: 6RK4) in complex with the AQKA stem peptide (Cα RMSD = 0.76 Å over 54 aligned residues). Host-adaptation residues in Lys1358 (S167, L186, and K187) and the corresponding positions in lysostaphin (T422, K441, and K442) are shown as sticks. In Lys1358, a co-crystallized acetate molecule (ACT) occupies a position that overlaps with the terminal D-Ala4 of the AEKA stem peptide in the lysostaphin complex; atoms are colored by element (C in grey/green/cyan, O in red, N in blue). **(b–d)** Electrostatic surface representations (scale from –5 to +5 k_BT/e) of (b) Lys1358 SH3b with bound acetate, (c) lysostaphin SH3b in complex with AQKA, and (d) a HADDOCK 2.4 docking model of the AQKA peptide bound to Lys1358. Ligands and docking models are highlighted in red. **(e)** Localized conformational plasticity in the L186–V192 region at the periphery of the stem peptide–binding cleft. Structural superposition of the high-resolution Lys1358 SH3b domain (1.22 Å) and the N66D variant shows that L186 adopts dual side-chain conformations (conformation A, green, 70% occupancy; conformation B, yellow, 30% occupancy). V192, located ∼3.5 Å from L186, adopts an alternative orientation in the N66D structure (magenta) relative to the WT.

Structural conservation of these SH3b domains, together with conservation of the stem peptide sequence between *L. lactis* and *S. aureus* ^3^, supports the use of lysostaphin as a framework to identify substrate-binding residues in Lys1358. In the Lys1358 crystal structure, a co-crystallized acetate molecule occupies a hydrophobic pocket that spatially overlaps with the terminal D-Ala4 of the AEKA stem peptide in the lysostaphin complex (Fig. 5a–c), suggesting that this pocket serves as a conserved recognition site for the stem-peptide C-terminus.

To validate this site, we mutated Arg172 (R172) and Trp203 (W203), which correspond to lysostaphin Arg427 (R427) and His458 (H458), respectively, reported to coordinate D-Ala4 ^25^. The R172A mutation significantly reduced activity, while the R172A/W203A double variant abolished it (Supplementary Fig. 9). These results confirm the essential role of this pocket in substrate anchoring and validate the use of lysostaphin as a structural model for Lys1358.

We next modeled the Lys1358–stem peptide complex using HADDOCK 2.4 ^31^. To ensure biological relevance, docking was guided by active-site restraints derived from residues observed in direct contact with the peptidoglycan in the lysostaphin–AQKA co-crystal structure (see Methods) ^25^. The docking simulations revealed a binding mode consistent with that observed in lysostaphin (Fig. 5d), enabling the evaluation of the host-adaptation mutation S167I.

In Lys1358, S167 corresponds to Thr422 in lysostaphin, a residue that contributes to interactions with the lysine of the stem peptide. The S167I substitution increased the predicted desolvation energy (–2.72 ± 0.73 vs. –2.13 ± 0.97 for WT), which is consistent with enhanced hydrophobic interactions. However, the overall docking score for the mutant was less favorable than that of the wild type (–74.62 ± 23.26 vs. –89.45 ± 45.28 arbitrary units; Supplementary Fig. 10). Notably, S167I also slightly increased thermal stability, with a Tm of 56°C compared to 55.5°C for the WT (Supplementary Fig. 8).

In contrast to S167I, the L186I mutation had minimal effects on docking energies and thermal stability. Furthermore, molecular dynamics (MD) simulations did not reveal significant large-scale deviations or stability shifts attributable to the L186I substitution. However, a high-resolution crystal structure (1.22 Å; PDB ID: 12LL) revealed local structural heterogeneity. In this structure, L186 is the only residue in the SH3b domain exhibiting dual side-chain conformations, adopting two distinct rotameric states with occupancies of 0.70 and 0.30, respectively.

This conformational plasticity extends to the immediate vicinity: V192 (Val192, ∼3.5 Å from L186) adopts an alternative orientation in the N66D variant structure relative to the wild-type protein. Because N66 is located in the distant catalytic domain, this structural divergence likely reflects intrinsic conformational sampling of the SH3b domain rather than a long-range allosteric effect triggered by the mutation. Together, the conformational heterogeneity at position 186 and the sampling of alternative V192 orientations suggest that this region, located at the periphery of the stem peptide binding cleft, may function as a localized flexible hinge.

### Compensatory mutation in a phage transglycosylase suggests adaptation to impaired endolysin function

During adaptive evolution, selective pressures act across the phage genome to optimize fitness, often through epistatic or compensatory interactions. To identify loci under selection beyond the engineered endolysin, we performed longitudinal whole-genome sequencing (Illumina) of wild-type phage P008 and the chimeric derivative P008::Lys1358 at passages P20, P30, and P40.

Comparative genomics identified R138W (Arg138Trp) as the only non-synonymous mutation reaching fixation specifically in the P008::Lys1358 lineage (Fig. 6a). This mutation occurs within *orf23*, a predicted transglycosylase located immediately downstream of the endolysin in the lysis module. Structural modelling using AlphaFold3 indicates that ORF23 comprises an N-terminal α-helical domain (residues 1–66) and a C-terminal lysozyme-like catalytic domain (residues 99–178) containing the R138W substitution. Structural homology searches identified hen egg-white lysozyme (HEWL; PDB ID: 1DPX) as a close match to this domain (Cα RMSD = 1.48 Å over 87 aligned residues; Fig. 6b).

**Fig. 6.**
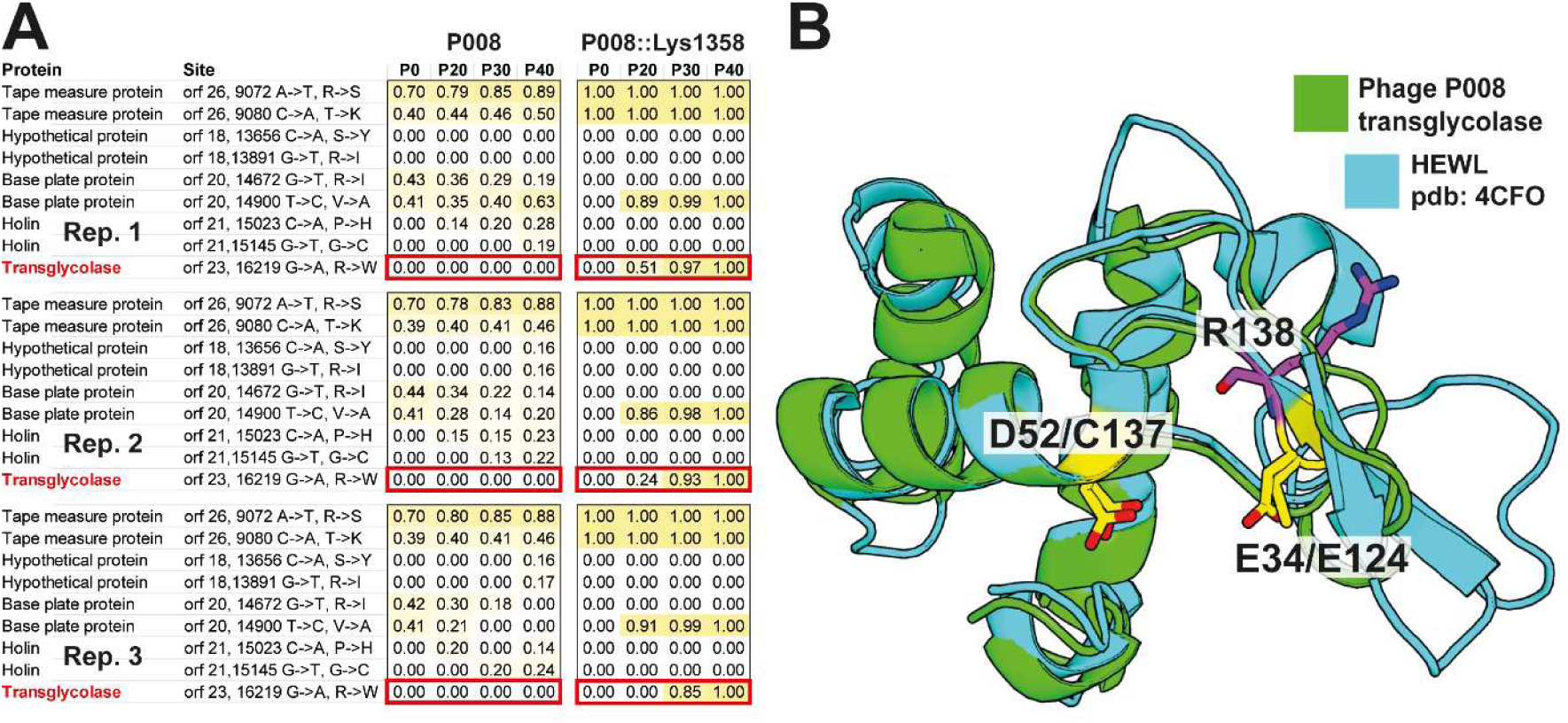
Adaptive genomic landscape and structural basis of compensatory evolution in ORF23. **(a)** Evolutionary dynamics of secondary mutations. Longitudinal tracking of non-synonymous mutation frequencies across the phage genome (excluding the endolysin locus) during serial passage (P0 to P40). The R138W mutation in *orf23* (transglycosylase) is uniquely fixed in the P008::Lys1358 lineage. **(b)** Structural conservation and active-site modulation of the ORF23 catalytic domain. Tertiary structure alignment of the predicted ORF23 C-terminal domain (residues 99–178, green) with hen egg-white lysozyme (HEWL, cyan; PDB ID: 1DPX). The catalytic dyad of HEWL (E35, D52) and corresponding putative active-site residues in ORF23 (E124, C137) are highlighted as yellow sticks. The adaptive R138W mutation (magenta) is positioned at the periphery of the active-site cleft, occupying a site analogous to Y53 in HEWL.

HEWL hydrolyzes the β-(1→4) glycosidic bond between NAM and NAG via a catalytic dyad composed of Glu35 (E35) and Asp52 (D52) ^32^. In ORF23, the catalytic glutamate is conserved as Glu124 (E124), while the position corresponding to D52 is occupied by Cys137 (C137). The spatial positioning of C137 suggests that it may serve as a functional analogue to D52 in HEWL, which is known to stabilize the transition state during catalysis ^32^. Although this represents a non-conservative substitution, both residues can nonetheless contribute to polar interactions and hydrogen-bond stabilization.

Adjacent to this catalytic center, the R138W mutation introduces a bulky aromatic residue at a position analogous to Tyr53 (Y53) in HEWL (Fig. 6b). Notably, tryptophan and tyrosine share similar physicochemical properties as large aromatic residues, suggesting convergence toward a structural feature characteristic of lysozyme active sites. Together, these observations are consistent with the hypothesis that ORF23 may be subject to compensatory adaptation under the altered peptidoglycan cleavage context imposed by the heterologous Lys1358 endolysin.

## Discussion

In this study, we investigate the molecular mechanisms underlying the evolution of a phage endolysin toward lysis of a novel bacterial host. Adaptation follows a dominant and highly reproducible trajectory, marked by the consistent fixation of three key mutations (N66D, S167I, and L186I), consistent with strong positive selection ^33^. Although these three substitutions were previously identified in our study of P008::Lys1358 and related chimeric phages ^14^, that work focused primarily on the evolutionary consequences of modular exchange. Restricted by short-read sequencing and the analysis of a single evolutionary trajectory, it lacked the resolution to determine mutation order, identify alternative pathways, or provide a mechanistic interpretation of the adaptive process.

Using PacBio deep sequencing across replicate evolutionary trajectories, we obtained full-length endolysin gene sequences from evolving phage populations, enabling tracking of allele frequency dynamics and inference of the sequential acquisition of mutations. N66D consistently fixed first, followed by S167I, which emerged concurrently but at lower frequency. The L186I mutation was never observed alone; instead, it appeared in combination with N66D, either as a double variant (N66D+L186I) or as part of the triple variant (N66D+S167I+L186I). The N66D+S167I double variant increased in frequency prior to the appearance of the triple variant, which emerged later and at lower frequencies, reflecting a deceleration of the adaptive process.

These reproducible evolutionary trajectories are a well-established feature of adaptive evolution and have been observed in both in vivo and in vitro model systems. For example, the evolution of an adenylate kinase toward increased thermostability ^34^ and a phage tail fiber protein toward a novel receptor-binding site ^35^ both follow predictable evolutionary paths. Likewise, in vitro studies on β-lactamase evolution, involving systematic combinations of clinically observed ^36^ or experimentally selected mutations ^37^, have shown that adaptation often proceeds along constrained mutational routes due to epistatic interactions. These constraints frequently impose a specific order of mutation acquisition and limit the number of accessible high-fitness variants.

In our study, however, the concurrent emergence of single variants (N66D or S167I) and double variants (N66D+S167I or N66D+L186I), albeit at different frequencies, suggests that negative epistasis did not impact their sequential acquisition. Instead, the differing frequencies of intermediate variants point to fitness differences, which were confirmed by biochemical assays showing that the order of fixation aligned with the relative gains in enzymatic activity conferred by each mutation. Importantly, no evidence of clonal interference was observed; mutations accumulated in a reproducible, stepwise manner, with little sign of competing lineages ^38^. This pattern reflects the classical dynamics of an adaptive walk under strong selection, where mutations of large effect tend to fix early due to their rapid spread and high fitness advantage ^39^. The observed deceleration toward the end of the trajectory is consistent with the principle of diminishing returns, where subsequent mutations yield progressively smaller fitness gains as the population nears a local fitness peak ^40,41^. While our results largely support a predictable and repeatable adaptive trajectory, one replicate deviated from this pattern, revealing a transient alternative evolutionary path that underscores the interplay between contingency and constraint. In this replicate, the canonical trajectory (N66D → N66D+S167I → N66D+S167I+L186I) was observed, but a parallel lineage also emerged: L186I+K187I appeared early and sequentially acquired N66D and S167I. This variant ultimately converged on a high-fitness genotype carrying the same three core mutations as the dominant trajectory, but with an additional K187I substitution. The rarity of this parallel path likely reflects the lower likelihood of initially acquiring the L186I+K187I combination compared to the more accessible and strongly beneficial N66D mutation. These findings highlight how adaptive walks, while often constrained and directional, can also accommodate occasional divergence due to rare events—reinforcing the view that evolution is shaped by both selection and contingency ^42–44^.

Our results illustrate a classic consequence of functional adaptation: the emergence of evolutionary trade-offs. Mutations that enhance performance in a new host often compromise activity in the ancestral context, reflecting antagonistic pleiotropy and functional specialization ^45,46^. The triple mutant (N66D, S167I, and L186I) identified at the end of the evolutionary trajectory displayed enhanced lytic activity on *L. lactis* IL1403, the host used during experimental evolution, but exhibited significantly reduced activity on *L. lactis* 582, the original host of phage 1358. This pattern is consistent with previous studies of enzyme specificity, where adaptation to a new substrate often reduces efficiency on the original one ^47^. These trade-offs extend to protein stability, as shown by differential scanning fluorimetry, which revealed that the first fixed mutation, N66D, is destabilizing. Collectively, these findings illustrate how adaptive mutations, while beneficial in terms of function, can incur structural costs, further emphasizing the complex fitness landscape navigated during protein evolution^29,48^.

Yet a fundamental question remains: how do adaptive mutations reshape the molecular architecture of a viral enzyme, initially characterized by suboptimal, promiscuous activity, to enhance function in a new host environment? Structural and mechanistic analyses revealed that the three adaptive mutations (N66D, S167I, and L186I) are located outside the catalytic site of the endolysin, indicating that the observed gains in lytic activity arise from improved substrate recognition rather than altered catalytic efficiency. ELISA assays confirmed that these mutations significantly increased binding affinity to the new host, with early-fixing variants, particularly N66D, showing the most pronounced gains.

The N66D substitution introduces a negative charge near a substrate-binding cleft, likely modifying the local electrostatic environment to enhance interactions with the peptidoglycan. Although this residue is located within a hand-cleft–like motif distal to the catalytic site, its position suggests a role in engaging an adjacent stem peptide repeat, consistent with the extended, polymeric architecture of the peptidoglycan matrix ^3^. Such modifications could expand the binding interface and increase avidity for the substrate. Similar electrostatic adjustments have been shown to modulate enzyme–substrate affinity by facilitating long-range interactions and stabilizing transition states ^49^. In phage endolysins, changes in surface charge have been linked to enhanced bacteriolytic activity through improved substrate interaction ^50^.

The other two mutations reside within the SH3b cell wall-binding domain and could be interpreted more precisely by leveraging its structural homology to lysostaphin’s SH3b domain, which was co-crystallized with a peptidoglycan stem peptide. Notably, the S167I substitution replaces a polar serine, capable of hydrogen bonding, with a bulky hydrophobic isoleucine at a site that, in lysostaphin, contacts a lysine side chain in the substrate. This change likely disrupts hydrogen bonding with the substrate while increasing hydrophobicity at the interface. The resulting decrease in desolvation energy reflects reduced energetic costs of water displacement, thereby strengthening binding via more favorable hydrophobic interactions ^51^. In parallel, this substitution conferred a modest increase in thermal stability, likely due to tighter packing and enhanced van der Waals contacts within local hydrophobic clusters ^52^.

While N66D and S167I impart clear physicochemical changes that directly enhance substrate engagement, L186I appears to act through subtle local conformational fine-tuning. High-resolution crystallography revealed that L186 adopts multiple rotameric states, indicating that this region at the periphery of the binding cleft behaves as a dynamic hinge. Substitution with the β-branched isoleucine likely restricts this mobility, promoting more defined packing interactions that stabilize the positioning of residues K187 and T191, a structural effect that has been associated with increased local stability in other protein systems ^53,54^. In the conserved lysostaphin SH3b binding interface, the corresponding residues (K442 and T446) exhibit significant chemical shift perturbations upon binding to the peptidoglycan stem peptide, implicating this region in substrate interaction ^25^. By modulating the dynamics of this region, L186I may optimize the orientation of these key contact points. The co-occurrence of L186I and K187I in an evolutionary replicate further suggests a permissive role for position 186; by stabilizing the local structural environment, L186I likely facilitates subsequent adaptive substitutions at the primary binding interface. Together, these results highlight an evolutionary strategy in which local structural tuning buffers the impact of mutations at key functional residues, expanding the accessible mutational landscape during protein evolution ^55,56^.

Such fine-tuning of enzyme activity may have been driven by subtle differences in cell wall architecture between *L. lactis* host strains. IL1403 exhibits a thinner wall and faster growth than *L. lactis* 582, suggesting reduced structural density and potentially looser crosslinking ^57^. These structural differences may provide a more favorable environment for the adaptive mutations (N66D, S167I, and L186I), which enhance substrate binding and improve lytic activity. For example, differences in crosslinking patterns have been shown to alter the activity of autolysins and phage-derived lytic enzymes by inhibiting substrate access and modulating enzymatic function ^58^.

Although this study focused on the endolysin gene, the entire phage genome was subject to mutational pressure during adaptation. Genome-wide analysis identified a single non-synonymous mutation, R138W, fixed exclusively in the P008::Lys1358 lineage, which occurs in a predicted lytic transglycosylase gene (orf23). Lytic transglycosylases are functional analogs of lysozymes that cleave the β-(1→4)-glycosidic bond between NAM and NAG residues via a conserved catalytic glutamate driving intramolecular cyclization rather than hydrolysis ^32,59^. Structural modelling revealed strong similarity between ORF23 and hen egg-white lysozyme (HEWL), with Glu124 acting as the catalytic proton donor and Cys137 aligning with Asp52 of HEWL, placing the R138W substitution near the predicted active site. The mutation introduces a bulky aromatic residue equivalent to Tyr53 in HEWL, potentially influencing the catalytic environment or substrate positioning by restoring an aromatic character typical of lysozyme active-site architectures.

We propose that this mutation reflects fine-tuning of ORF23 activity in response to altered peptidoglycan processing resulting from replacement of the ancestral amidase 2 activity by the CHAP–SH3 endolysin. Given that ORF23 is located downstream of the endolysin gene, this may facilitate functional coordination between enzymes acting on the same substrate pool. Such cooperative interactions are a well-established feature of phage lysis systems, where multiple enzymes act synergistically to breach the bacterial cell wall, consistent with our findings that adaptation in the lysis module can involve compensatory mutations alongside primary effectors to achieve integrated fitness gains ^60,61^.

Taken together, our study reveals that phage endolysins evolve through reproducible, stepwise accumulation of mutations that fine-tune substrate recognition and enzymatic performance in a new host context. This adaptive process reflects a balance between structural constraints, selection, and occasional contingency, resulting in functionally specialized variants optimized for distinct bacterial cell wall architectures. Importantly, the predictability of these adaptive trajectories suggests that evolutionary dynamics can provide a framework for guiding the engineering of endolysins for application-specific functions, including targeting multidrug-resistant bacteria and tailoring activity to defined environmental contexts.

## Acknowledgments

X.Z. was supported by graduate scholarships from the Fonds de recherche du Québec – Nature et technologies. V.S. was supported by a postdoctoral fellowship from the Swiss National Science Foundation under grant P500PB_214419. R.A.C. acknowledges grants from the Natural Sciences and Engineering Research Council of Canada (RGPIN-2021-03484 and RGPAS-2021-00017) and the Canada Foundation for Innovation (26503). This research was enabled in part by support provided by Compute Ontario (www.computeontario.ca) and the Digital Research Alliance of Canada (alliancecan.ca). R.S. and S.M. are grateful to PROTEO for funding through its New Initiative Program. The crystallographic work was partially supported by an NSERC Discovery Grant (RGPIN-2020-06954) to R.S. S.M. also acknowledges funding from the Natural Sciences and Engineering Research Council of Canada (NSERC; Discovery and Alliance programs) and holds a Tier 1 Canada Research Chair in Bacteriophages. F.O. was supported by a postdoctoral fellowship from the Swiss National Science Foundation (P400PB_191059) and by an NSERC Discovery Grant (RGPIN-2025-05623).

This research used resources of the Advanced Photon Source; a U.S. Department of Energy (DOE) Office of Science User Facility operated for the DOE Office of Science by Argonne National Laboratory under Contract No. DE-AC02-06CH11357. Use of the Lilly Research Laboratories Collaborative Access Team (LRL-CAT) beamline at Sector 31 of the Advanced Photon Source was provided by Eli Lilly and Company, which operates the facility. Part of the research described in this paper was performed using beamline CMCF-BM at the Canadian Light Source, a national research facility of the University of Saskatchewan, which is supported by the Canada Foundation for Innovation (CFI), the Natural Sciences and Engineering Research Council (NSERC), the National Research Council (NRC), the Canadian Institutes of Health Research (CIHR), the Government of Saskatchewan, and the University of Saskatchewan.

## Contributions

F.O. conceived the study, led the project, and performed the majority of the experimental work. F.O. analysed the data, performed structural and docking analyses, prepared Figs. 1, 3–6, and wrote the original manuscript. X.Z. contributed to protein purification and, with R.S., determined the crystal structures of wild-type Lys1358 and the N66D mutant. V.S. analysed PacBio and Illumina sequencing data and generated Fig. 2. R.V.R. and R.A.C. performed molecular dynamics simulations. F.O. and S.M. supervised the study and contributed to experimental design and data interpretation. All authors contributed to data interpretation, discussed the results, and revised the manuscript.

## Methods

### Bacterial strains, phages, and growth conditions

The biological materials and plasmids used in this study are listed in Supplementary Table 8. *L. lactis* strains were grown without agitation at 30 °C in M17 broth supplemented with 0.5% glucose (GM17) and 1.0% agar (for solid media). *E. coli* strains were grown at 37 °C with agitation (220 rpm) in LB or plated on LB with agar (LBA). If necessary, LB was supplemented with kanamycin sulfate (50 μg/mL). For phage infection, 10 mM CaCl_2_ was added to GM17, and the double-layer plaque assays were performed as previously described [1].

### Experimental evolution of phages P008 and P008::Lys1358 and sequencing analysis

The Lactococcal phage P008 and the endolysin variant P008::Lys1358 were evolved over 40 serial transfers in triplicate. *L. lactis* IL1403 was grown in 10 mL volumes in test tubes to early exponential phase (OD₆₀₀ ∼0.2). For the initial infection, 100 µL of a 10⁷ pfu/mL phage stock was added to the cultures, which were then incubated overnight at 30 °C. Lysed cultures were subsequently filtered through a 0.4 µm filter and diluted 100-fold. For each subsequent transfer, 100 µL of this diluted lysate was used to infect fresh bacterial cultures, which were again incubated overnight at 30 °C. Lytic plaque surface areas were measured between passages using image analysis, as previously described [2].

### Genomic sequencing and analysis of evolved phage populations

For PacBio sequencing of the *Lys1358* gene during experimental evolution, phage genomic DNA was extracted from each triplicate every two transfers, following the procedure described previously (Moineau et al., 1994). The *Lys1358* gene was amplified from each condition using primers with unique barcodes for each passage and replicate, a high-fidelity polymerase (KAPA HiFi, Roche), and approximately 2 ng of DNA per PCR reaction. PCR product size and quantity were verified by agarose gel electrophoresis, and equal volumes of each barcoded amplicon were pooled in a 2.0 mL DNA LoBind tube for a total volume of 800 µL. Multiplexed amplicon libraries were prepared using the SMRTbell Prep Kit 3.0 and sequenced on a Revio 25M flow cell at SeqCenter LLC, PA, USA.

Adapter sequences were removed using Cutadapt ^62^. Reads were mapped to the amplicon reference using Minimap2 ^63^, and single nucleotide polymorphisms (SNPs) were identified with default parameters of FreeBayes ^64^. Based on the observed error rate in the isogenic starting population (P₀), a 1% allele frequency threshold was applied to define reliable de novo mutations. To assess linkage (synteny) among mutations, individual reads were screened for the presence of reference and alternative alleles using custom grep-based parsing. Data processing and visualization were performed in R ^65^, with figures generated using ggplot2 ^66^. Growth curves were fitted using the growthcurver ^67^, and Muller plots were generated with ggmuller ^68^.

For whole-genome sequencing of phage populations using Illumina technology, libraries were prepared with the Nextera XT DNA Library Preparation Kit (Illumina), following the manufacturer’s instructions. Sequencing was performed on a MiSeq system with a MiSeq Reagent Kit v2 (Illumina). Illumina reads were adapter- and quality-trimmed using Trim Galore (DOI: 10.5281/zenodo.5127899) and mapped to the reference genome (P008_Lys1358) using BWA-MEM ^69^. Single nucleotide polymorphisms (SNPs) were called using FreeBayes with parameters -C 5 --pooled-continuous --min-alternate-fraction 0.05 --min-coverage 10.

### Cloning, expression, and purification of Lys1358 and its variant derivatives

Plasmids were purified from overnight *E. coli* cultures using the QIAprep Spin Miniprep Kit (Qiagen). Polymerase chain reactions (PCR) were carried out with Taq polymerase (Feldan) for screening or with Q5 High-Fidelity DNA Polymerase (New England Biolabs) for site-directed mutagenesis. The plasmids and primers used in this study are listed in Supplementary Table 8.

The pLys1358a construct, based on the pET28a vector encoding Lys1358 with a C-terminal His-tag [2], served as the template for mutagenesis. Amino acid substitutions identified during experimental evolution (N66D, S167I, L186I) were introduced, as well as alanine substitutions to generate inactive variants (C29A, R172S, W203A). Mutagenesis was performed using the Q5® Site-Directed Mutagenesis Kit (NEB) with primer pairs designed using the manufacturer’s online tool (nebasechanger.neb.com). All constructs were verified by sequencing. Lys1358 and its variants were expressed and purified by nickel affinity chromatography as previously described [2].

### Evaluation of the activity of Lys1358 and derivatives on *L. lactis* cells

The lytic activity of Lys1358 and its variant derivatives were evaluated by monitoring the reduction in turbidity of a solution containing *L. lactis* IL1403 or 582 cells at an OD_600nm_ of 0.4. Bacterial cells were resuspended in lysis buffer (20 mM Tris-HCL, 200 mM NaCl, 2mM CaCl2, pH 7.0) and mixed in a 96-well microplate with purified endolysin diluted to the desired concentration in endolysin buffer (20 mM Tris pH 7.3, 150 mM NaCl, 2mM CaCl2). The decrease in turbidity was measured at 37 °C using a microplate reader (BioTek Synergy HTX Multimode Reader).

### ELISA assessment of endolysin binding to bacterial cells

*L. lactis* IL1403 and 582 cells were grown to an OD₆₀₀ of ∼0.4, harvested, washed twice with PBS, and concentrated 10-fold. Cells (200 µL/well) were immobilized on Nunc Immunoplates (MaxiSorp, Sigma-Aldrich, M5785-1CS) overnight at 4 °C. Wells were washed four times with PBS and blocked with 3% BSA in PBS for 1 h at room temperature. catalytically inactive endolysin variants Endolysins were prepared as 10-fold serial dilutions in PBS, added to the wells (100 µL), and incubated for 1 h at room temperature. After three PBS washes, bound protein was detected with 100 µL anti-His primary antibody (1:1000 in PBS + 0.01% Tween-20; MP Biomedicals™, MP08L100033) for 1 h, followed by three washes and incubation with 100 µL secondary antibody (1:2000 in PBS + 3% BSA; Goat anti-Rabbit IgG HRP, Invitrogen 31460) for 1 h. Color was developed with 100 µL TMB substrate (Thermo Scientific PI34021) and stopped with 100 µL 1 M H₂SO₄. Absorbance was measured at 450 nm using a BioTek Synergy HTX Reader.

### Crystallization, data collection, and structure determination

Crystallization of Lys1358 and its N66D variant after surface lysine methylation was performed at room temperature via the micro-batch-under-oil approach, where the protein and reservoir solution were mixed at a 1:1 ratio (v/v). The Lys1358 crystals were obtained using the reservoir solution containing 0.1 M sodium acetate pH 4.5 and 2.0 M ammonium sulfate while the N66D variant was crystallized in the solution containing 0.2 M ammonium sulfate, 0.1 M sodium acetate pH 4.6, and 30% PEG 2000 MME. The Lys1358 crystals is in the space group C2 with the unit cell parameters a=123.7, b=39.5, c=56.1 Å, β=108.8° while the N66D variant is in the I2 space group with a=55.7, b=39.3, c=118.4 Å, β=97.5°. The crystals were cryoprotected by the reservoir solution supplemented with 30% glycerol before data collection at the LRL-CAT (31-ID) beamline at Advanced Photon Source, Argonne National Laboratory or at the CMCF-BM beamline at Canadian Light Source. Data processing and scaling were performed with XDS ^70^. The initial phase of Lys1358 structure was determined by MolRep ^71^ using the CHAP domain (res 1-143) of the endolysin LysIME-EF1 (PDB 6IST) and was then completed via multiple cycles of ARP/wARP model building ^72^. The structure of the N66D variant was determined by MolRep using the mLys1358 model as the template. To get the final structures, multiple cycles of refinement using Refmac5 ^73^ in CCP4 ^74^ followed by manual model rebuilding with Coot ^75^ were carried out. The stereochemistry of the final model has been analyzed with PROCHECK ^76^. Data collection and refinement statistics are shown in Supplementary Table 7. The coordinates and structure factors of Lys1358 and Lys1358 N66D variant have been deposited in the Protein Data Bank with the accession codes 10VD, 12LL, and 10VE, respectively.

### Molecular dynamics simulations

All molecular dynamics (MD) simulations were performed using AMBER 2020 (http://ambermd.org/) with the AMBER19SB forcefield^77^. The crystal structure of wild-type Lys1358 was used as the starting model. Structures of the N66D, N66D/S167I, and N66D/S167I/L186I variants were generated *in silico* from the wild-type crystal structure using the protein design software Triad (Protabit LLC, Pasadena, CA, USA) ^78^. Amino-acid protonation states were predicted using the H + + server (http://newbiophysics.cs.vt.edu/H++/). Each enzyme structure was solvated in a pre-equilibrated orthorhombic box of TIP3P water extending 8 Å beyond the solute in all directions and neutralized with explicit Na⁺ and Cl⁻ counterions using the LEaP module ^79^. Energy minimization was performed in two stages: (i) solvent and ion minimization with harmonic positional restraints on the solute (500 kcal mol⁻¹ Å⁻²), followed by (ii) unrestrained minimization of all atoms. The systems were gradually heated from 0 K to 300 K in six 50 ps steps (ΔT = 50 K) under constant volume and periodic boundary conditions. Water bonds were constrained using the SHAKE algorithm, and long-range electrostatics were treated with the particle-mesh Ewald method using an 8 Å cutoff for nonbonded interactions ^79^. Harmonic restraints on the protein were progressively reduced (210, 165, 125, 85, 45, 10 kcal mol⁻¹ Å⁻²) during heating, with temperature control maintained using the Langevin thermostat. A 2 fs timestep was used throughout. Following heating, each system was equilibrated for 20 ns in the NPT ensemble (1 atm, 300 K) without restraints, followed by 1 μs production simulations in the NVT ensemble with periodic boundary conditions. Three independent production replicates were performed for each variant.

### Differential scanning fluorimetry

Solutions of Lys1358 and variant variants (N66D, S167I, L186I, and the triple variant N66D/S167I/L186I) were prepared at 1 mg·mL⁻¹ in the protein storage buffer. For each reaction, 20 µL of protein solution was mixed with 5 µL of JBS Thermofluor dye (Jena Bioscience) diluted 1:500 in the same buffer. Following a 2-min equilibration at 4 °C, samples were heated from 4 °C to 95 °C at 0.5 °C·min⁻¹ in a CFX96 Touch Real-Time PCR Detection System (Bio-Rad). Fluorescence was recorded in the FRET channel throughout the temperature ramp and data were processed using Bio-Rad CFX Maestro. Melting temperatures (Tₘ) were determined from the minimum of the first derivative of the fluorescence signal.

### Docking Prediction of the Lys1358 SH3b Domain with the Stem Peptide AQKA

The structure of the Lys1358 SH3b domain was predicted using AlphaFold via the AlphaFold server (https://alphafoldserver.com/) for the wild-type (WT) and variants S167L, L187I, and the double variant S167L/L187I. The AQKA stem peptide was extracted from the crystal structure of the lysostaphin SH3b domain (PDB ID: 6RK4), where it was originally co-crystallized. Docking simulations were performed using HADDOCK 2.4 with default parameters ^31^. Residues previously reported to interact with the stem peptide in lysostaphin by NMR and crystallography (NmR: N421/N166, I425/N170, T426/I171, S438/A183, V440/V185, K442/K187, A443/A188, V461/V206; Structure: T422, S167, D423/A168, I424/V169, I425/N170, R427/R172, R433/T177, H458/W203, W460/W205, P474/A219, W489/T229) were defined as active residues for the docking ^25^. For each structure, 200 complexes were generated and manually curated by visual alignment of the peptide to its position in the lysostaphin reference structure. Only complexes in which the peptide adopted a similar orientation and position were retained for energy evaluation (see Supplementary Fig. 12 for an example).

## Statistical analysis

Statistical analyses were performed in R (v4.x). Data are reported as mean ± standard deviation (s.d.). Differences in cell wall thickness between *L. lactis* IL1403 and *L. lactis* 582 were assessed using a two-sided Welch’s t-test. Plaque size data are presented as mean ± s.d. For comparisons between wild-type P008 and P008::Lys1358 at passage 0 (P0), normality was assessed using the Shapiro–Wilk test. As data deviated from normality, differences between groups were evaluated using a two-sided Wilcoxon rank-sum test. Changes in plaque size over experimental evolution were quantified by linear regression, treating passage number as a continuous variable. The significance of the regression slope was assessed using a two-sided t-test. Lytic activity and ELISA binding data were analysed using linear mixed-effects models, with construct (WT and variants), enzyme concentration, and their interaction included as fixed effects, and biological replicate as a random effect. Pairwise comparisons between constructs at each concentration were performed using estimated marginal means (EMMs). P values were adjusted within each concentration where applicable. Statistical significance was defined as P < 0.05.

