## Supplementary Tables for "Predictable adaptive evolution of a phage endolysin through substrate recognition optimization"

**Supplementary Table 1.** Estimated differences in lytic activity of endolysin variants relative to Lys1358 across enzyme concentrations. Values were obtained from linear mixed-effects models followed by estimated marginal means (EMMs) contrasts at each enzyme concentration (µM). All data were obtained using *L. lactis* IL1403. Positive values indicate higher activity relative to Lys1358. The "Triple" construct corresponds to the combined N66D, S167I, and L186I substitutions. SE, standard error; df, degrees of freedom; t ratio, test statistic from model contrasts; P value, significance of the comparison. Statistical significance is indicated as follows: *P < 0.05; **P < 0.01; ***P < 0.001.

| Concentration (µM) | Construct | Estimate vs WT | SE | df | t ratio | P value | Significance |
|---|---|---|---|---|---|---|---|
| 0.05 | L186I | 2.686215 | 1.701379 | 239 | 1.578845 | 0.329 | |
| 0.05 | N66D | 2.8414505 | 1.701379 | 239 | 1.670086 | 0.282 | |
| 0.05 | S167I | 2.8901139 | 1.701379 | 239 | 1.698689 | 0.269 | |
| 0.05 | Triple | 45.8684619 | 1.701379 | 239 | 26.95957 | < 0.0001 | *** |
| 0.08 | L186I | 4.3170108 | 1.701379 | 239 | 2.537359 | 0.041 | * |
| 0.08 | N66D | 15.9625716 | 1.701379 | 239 | 9.382136 | < 0.0001 | *** |
| 0.08 | S167I | 4.5812306 | 1.701379 | 239 | 2.692657 | 0.027 | * |
| 0.08 | Triple | 60.4354679 | 1.701379 | 239 | 35.52145 | < 0.0001 | *** |
| 0.1 | L186I | 13.219431 | 1.701379 | 239 | 7.769832 | < 0.0001 | *** |
| 0.1 | N66D | 37.2719376 | 1.701379 | 239 | 21.90689 | < 0.0001 | *** |
| 0.1 | S167I | 15.4285822 | 1.701379 | 239 | 9.068279 | < 0.0001 | *** |
| 0.1 | Triple | 68.4273957 | 1.701379 | 239 | 40.21878 | < 0.0001 | *** |
| 0.15 | L186I | 26.4978317 | 1.701379 | 239 | 15.57432 | < 0.0001 | *** |
| 0.15 | N66D | 55.2225946 | 1.701379 | 239 | 32.45754 | < 0.0001 | *** |
| 0.15 | S167I | 41.1454659 | 1.701379 | 239 | 24.18359 | < 0.0001 | *** |
| 0.15 | Triple | 61.9153897 | 1.701379 | 239 | 36.39129 | < 0.0001 | *** |
| 0.2 | L186I | 4.3746881 | 1.701379 | 239 | 2.57126 | 0.038 | * |
| 0.2 | N66D | 12.8027234 | 1.701379 | 239 | 7.524908 | < 0.0001 | *** |
| 0.2 | S167I | 10.4554183 | 1.701379 | 239 | 6.14526 | < 0.0001 | *** |
| 0.2 | Triple | 14.9877119 | 1.701379 | 239 | 8.809154 | < 0.0001 | *** |
| 0.3 | L186I | -2.4791299 | 1.701379 | 239 | -1.45713 | 0.397 | |

| | | | | | | |
|---|---|---|---|---|---|---|
| 0.3 | N66D | 1.3136878 | 1.701379 | 239 | 0.772131 | 0.816 |
| 0.3 | S167I | 2.2188707 | 1.701379 | 239 | 1.30416 | 0.490 |
| 0.3 | Triple | 3.473666 | 1.701379 | 239 | 2.041676 | 0.138 |
| 0.45 | L186I | -0.287517 | 1.701379 | 239 | -0.16899 | 0.995 |
| 0.45 | N66D | 0.1785424 | 1.701379 | 239 | 0.10494 | 0.998 |
| 0.45 | S167I | 1.2155697 | 1.701379 | 239 | 0.714461 | 0.845 |
| 0.45 | Triple | 1.7504835 | 1.701379 | 239 | 1.028861 | 0.66 |
| 0.6 | L186I | -1.168017 | 1.701379 | 239 | -0.68651 | 0.858 |
| 0.6 | N66D | -0.9708564 | 1.701379 | 239 | -0.57063 | 0.907 |
| 0.6 | S167I | 0.5466674 | 1.701379 | 239 | 0.321308 | 0.976 |
| 0.6 | Triple | 1.7189061 | 1.701379 | 239 | 1.010302 | 0.678 |

**Supplementary Table 2.** Estimated marginal means (EMMs) of lytic activity for Lys1358 and variants at each enzyme concentration (μM), derived from linear mixed-effects models. All data were obtained using *L. lactis* IL1403. The "Triple" construct corresponds to the combined N66D, S167I, and L186I substitutions. WT corresponds to the unmutated Lys1358 enzyme. SE, standard error; df, degrees of freedom; Lower CL, lower confidence limit; Upper CL, upper confidence limit. Statistical significance is indicated as follows: *P < 0.05; **P < 0.01; ***P < 0.001.

| Concentration (μM) | Construct | Marginal mean | SE | df | Lower CL | Upper CL |
|---|---|---|---|---|---|---|
| 0.05 | L186I | 3.382 | 1.204315 | 215.18 | 1.00823 | 5.75576 |
| 0.05 | N66D | 3.53723 | 1.204315 | 215.18 | 1.16347 | 5.911 |
| 0.05 | S167I | 3.58589 | 1.204315 | 215.18 | 1.21213 | 5.95966 |
| 0.05 | Triple | 46.56424 | 1.204315 | 215.18 | 44.19048 | 48.93801 |
| 0.05 | WT | 0.69578 | 1.204315 | 215.18 | -1.67798 | 3.06954 |
| 0.08 | L186I | 6.93443 | 1.204315 | 215.18 | 4.56067 | 9.30819 |
| 0.08 | N66D | 18.57999 | 1.204315 | 215.18 | 16.20623 | 20.95375 |
| 0.08 | S167I | 7.19865 | 1.204315 | 215.18 | 4.82489 | 9.57241 |
| 0.08 | Triple | 63.05289 | 1.204315 | 215.18 | 60.67912 | 65.42665 |
| 0.08 | WT | 2.61742 | 1.204315 | 215.18 | 0.24366 | 4.99118 |
| 0.1 | L186I | 20.46057 | 1.204315 | 215.18 | 18.08681 | 22.83433 |
| 0.1 | N66D | 44.51308 | 1.204315 | 215.18 | 42.13931 | 46.88684 |
| 0.1 | S167I | 22.66972 | 1.204315 | 215.18 | 20.29596 | 25.04349 |
| 0.1 | Triple | 75.66854 | 1.204315 | 215.18 | 73.29477 | 78.0423 |
| 0.1 | WT | 7.24114 | 1.204315 | 215.18 | 4.86738 | 9.6149 |
| 0.15 | L186I | 46.40747 | 1.204315 | 215.18 | 44.03371 | 48.78124 |
| 0.15 | N66D | 75.13223 | 1.204315 | 215.18 | 72.75847 | 77.506 |
| 0.15 | S167I | 61.05511 | 1.204315 | 215.18 | 58.68134 | 63.42887 |
| 0.15 | Triple | 81.82503 | 1.204315 | 215.18 | 79.45127 | 84.19879 |
| 0.15 | WT | 19.90964 | 1.204315 | 215.18 | 17.53588 | 22.2834 |
| 0.2 | L186I | 73.48322 | 1.204315 | 215.18 | 71.10946 | 75.85699 |
| 0.2 | N66D | 81.91126 | 1.204315 | 215.18 | 79.5375 | 84.28502 |

| | | | | | | |
|---|---|---|---|---|---|---|
| 0.2 | S167I | 79.56395 | 1.204315 | 215.18 | 77.19019 | 81.93772 |
| 0.2 | Triple | 84.09625 | 1.204315 | 215.18 | 81.72248 | 86.47001 |
| 0.2 | WT | 69.10854 | 1.204315 | 215.18 | 66.73477 | 71.4823 |
| 0.3 | L186I | 81.36745 | 1.204315 | 215.18 | 78.99369 | 83.74122 |
| 0.3 | N66D | 85.16027 | 1.204315 | 215.18 | 82.78651 | 87.53403 |
| 0.3 | S167I | 86.06545 | 1.204315 | 215.18 | 83.69169 | 88.43922 |
| 0.3 | Triple | 87.32025 | 1.204315 | 215.18 | 84.94648 | 89.69401 |
| 0.3 | WT | 83.84658 | 1.204315 | 215.18 | 81.47282 | 86.22035 |
| 0.45 | L186I | 85.02441 | 1.204315 | 215.18 | 82.65064 | 87.39817 |
| 0.45 | N66D | 85.49047 | 1.204315 | 215.18 | 83.1167 | 87.86423 |
| 0.45 | S167I | 86.52749 | 1.204315 | 215.18 | 84.15373 | 88.90126 |
| 0.45 | Triple | 87.06241 | 1.204315 | 215.18 | 84.68864 | 89.43617 |
| 0.45 | WT | 85.31192 | 1.204315 | 215.18 | 82.93816 | 87.68569 |
| 0.6 | L186I | 83.72082 | 1.204315 | 215.18 | 81.34705 | 86.09458 |
| 0.6 | N66D | 83.91798 | 1.204315 | 215.18 | 81.54421 | 86.29174 |
| 0.6 | S167I | 85.4355 | 1.204315 | 215.18 | 83.06174 | 87.80926 |
| 0.6 | Triple | 86.60774 | 1.204315 | 215.18 | 84.23398 | 88.9815 |
| 0.6 | WT | 84.88883 | 1.204315 | 215.18 | 82.51507 | 87.2626 |

**Supplementary Table 3.** Pairwise comparisons of Lys1358 and variants lytic activity across enzyme concentrations. Estimated differences in lytic activity between constructs were obtained from estimated marginal means (EMMs)-based contrasts of a linear mixed-effects model at each enzyme concentration (µM). All data were obtained using *L. lactis* IL1403. The "Triple" construct corresponds to the combined N66D, S167I, and L186I substitutions. SE, standard error; df, degrees of freedom; t ratio, test statistic from model contrasts; P value, significance of the comparison. Statistical significance is indicated as follows: *P < 0.05; **P < 0.01; ***P < 0.001.

| Concentration (µM) | Contrast | Estimate | SE | df | t ratio | P value | Significance |
|---|---|---|---|---|---|---|---|
| 0.05 | L186I - N66D | -0.1552 | 1.7 | 239 | -0.091 | 1 | |
| 0.05 | L186I - S167I | -0.2039 | 1.7 | 239 | -0.12 | 1 | |
| 0.05 | L186I - Triple | -43.1823 | 1.7 | 239 | -25.381 | <0.0001 | *** |
| 0.05 | L186I - WT | 2.6862 | 1.7 | 239 | 1.579 | 0.5124 | |
| 0.05 | N66D - S167I | -0.0487 | 1.7 | 239 | -0.029 | 1 | |
| 0.05 | N66D - Triple | -43.027 | 1.7 | 239 | -25.289 | <0.0001 | *** |
| 0.05 | N66D - WT | 2.8415 | 1.7 | 239 | 1.67 | 0.4545 | |
| 0.05 | S167I - Triple | -42.9783 | 1.7 | 239 | -25.261 | <0.0001 | *** |
| 0.05 | S167I - WT | 2.8901 | 1.7 | 239 | 1.699 | 0.4367 | |
| 0.05 | Triple - WT | 45.8685 | 1.7 | 239 | 26.96 | <0.0001 | *** |
| 0.08 | L186I - N66D | -11.6456 | 1.7 | 239 | -6.845 | <0.0001 | *** |
| 0.08 | L186I - S167I | -0.2642 | 1.7 | 239 | -0.155 | 0.9999 | |
| 0.08 | L186I - Triple | -56.1185 | 1.7 | 239 | -32.984 | <0.0001 | *** |
| 0.08 | L186I - WT | 4.317 | 1.7 | 239 | 2.537 | 0.0859 | |
| 0.08 | N66D - S167I | 11.3813 | 1.7 | 239 | 6.689 | <0.0001 | *** |
| 0.08 | N66D - Triple | -44.4729 | 1.7 | 239 | -26.139 | <0.0001 | *** |
| 0.08 | N66D - WT | 15.9626 | 1.7 | 239 | 9.382 | <0.0001 | *** |
| 0.08 | S167I - Triple | -55.8542 | 1.7 | 239 | -32.829 | <0.0001 | *** |
| 0.08 | S167I - WT | 4.5812 | 1.7 | 239 | 2.693 | 0.058 | |
| 0.08 | Triple - WT | 60.4355 | 1.7 | 239 | 35.521 | <0.0001 | *** |
| 0.1 | L186I - N66D | -24.0525 | 1.7 | 239 | -14.137 | <0.0001 | *** |

| | | | | | | | |
|---|---|---|---|---|---|---|---|
| 0.1 | L186I - S167I | -2.2092 | 1.7 | 239 | -1.298 | 0.6924 | |
| 0.1 | L186I - Triple | -55.208 | 1.7 | 239 | -32.449 | <0.0001 | *** |
| 0.1 | L186I - WT | 13.2194 | 1.7 | 239 | 7.77 | <0.0001 | *** |
| 0.1 | N66D - S167I | 21.8434 | 1.7 | 239 | 12.839 | <0.0001 | *** |
| 0.1 | N66D - Triple | -31.1555 | 1.7 | 239 | -18.312 | <0.0001 | *** |
| 0.1 | N66D - WT | 37.2719 | 1.7 | 239 | 21.907 | <0.0001 | *** |
| 0.1 | S167I - Triple | -52.9988 | 1.7 | 239 | -31.151 | <0.0001 | *** |
| 0.1 | S167I - WT | 15.4286 | 1.7 | 239 | 9.068 | <0.0001 | *** |
| 0.1 | Triple - WT | 68.4274 | 1.7 | 239 | 40.219 | <0.0001 | *** |
| 0.15 | L186I - N66D | -28.7248 | 1.7 | 239 | -16.883 | <0.0001 | *** |
| 0.15 | L186I - S167I | -14.6476 | 1.7 | 239 | -8.609 | <0.0001 | *** |
| 0.15 | L186I - Triple | -35.4176 | 1.7 | 239 | -20.817 | <0.0001 | *** |
| 0.15 | L186I - WT | 26.4978 | 1.7 | 239 | 15.574 | <0.0001 | *** |
| 0.15 | N66D - S167I | 14.0771 | 1.7 | 239 | 8.274 | <0.0001 | *** |
| 0.15 | N66D - Triple | -6.6928 | 1.7 | 239 | -3.934 | 0.001 | *** |
| 0.15 | N66D - WT | 55.2226 | 1.7 | 239 | 32.458 | <0.0001 | *** |
| 0.15 | S167I - Triple | -20.7699 | 1.7 | 239 | -12.208 | <0.0001 | *** |
| 0.15 | S167I - WT | 41.1455 | 1.7 | 239 | 24.184 | <0.0001 | *** |
| 0.15 | Triple - WT | 61.9154 | 1.7 | 239 | 36.391 | <0.0001 | *** |
| 0.2 | L186I - N66D | -8.428 | 1.7 | 239 | -4.954 | <0.0001 | *** |
| 0.2 | L186I - S167I | -6.0807 | 1.7 | 239 | -3.574 | 0.0039 | * |
| 0.2 | L186I - Triple | -10.613 | 1.7 | 239 | -6.238 | <0.0001 | *** |
| 0.2 | L186I - WT | 4.3747 | 1.7 | 239 | 2.571 | 0.079 | |
| 0.2 | N66D - S167I | 2.3473 | 1.7 | 239 | 1.38 | 0.6413 | |
| 0.2 | N66D - Triple | -2.185 | 1.7 | 239 | -1.284 | 0.7012 | |
| 0.2 | N66D - WT | 12.8027 | 1.7 | 239 | 7.525 | <0.0001 | *** |
| 0.2 | S167I - Triple | -4.5323 | 1.7 | 239 | -2.664 | 0.0625 | |
| 0.2 | S167I - WT | 10.4554 | 1.7 | 239 | 6.145 | <0.0001 | *** |
| 0.2 | Triple - WT | 14.9877 | 1.7 | 239 | 8.809 | <0.0001 | *** |
| 0.3 | L186I - N66D | -3.7928 | 1.7 | 239 | -2.229 | 0.1725 | |
| 0.3 | L186I - S167I | -4.698 | 1.7 | 239 | -2.761 | 0.0483 | |

| | | | | | | |
|---|---|---|---|---|---|---|
| 0.3 | L186I - Triple | -5.9528 | 1.7 | 239 | -3.499 | 0.005 |
| 0.3 | L186I - WT | -2.4791 | 1.7 | 239 | -1.457 | 0.5913 |
| 0.3 | N66D - S167I | -0.9052 | 1.7 | 239 | -0.532 | 0.984 |
| 0.3 | N66D - Triple | -2.16 | 1.7 | 239 | -1.27 | 0.7102 |
| 0.3 | N66D - WT | 1.3137 | 1.7 | 239 | 0.772 | 0.9384 |
| 0.3 | S167I - Triple | -1.2548 | 1.7 | 239 | -0.738 | 0.9475 |
| 0.3 | S167I - WT | 2.2189 | 1.7 | 239 | 1.304 | 0.6889 |
| 0.3 | Triple - WT | 3.4737 | 1.7 | 239 | 2.042 | 0.2494 |
| 0.45 | L186I - N66D | -0.4661 | 1.7 | 239 | -0.274 | 0.9988 |
| 0.45 | L186I - S167I | -1.5031 | 1.7 | 239 | -0.883 | 0.9028 |
| 0.45 | L186I - Triple | -2.038 | 1.7 | 239 | -1.198 | 0.7526 |
| 0.45 | L186I - WT | -0.2875 | 1.7 | 239 | -0.169 | 0.9998 |
| 0.45 | N66D - S167I | -1.037 | 1.7 | 239 | -0.61 | 0.9735 |
| 0.45 | N66D - Triple | -1.5719 | 1.7 | 239 | -0.924 | 0.8875 |
| 0.45 | N66D - WT | 0.1785 | 1.7 | 239 | 0.105 | 1 |
| 0.45 | S167I - Triple | -0.5349 | 1.7 | 239 | -0.314 | 0.9979 |
| 0.45 | S167I - WT | 1.2156 | 1.7 | 239 | 0.714 | 0.953 |
| 0.45 | Triple - WT | 1.7505 | 1.7 | 239 | 1.029 | 0.8418 |
| 0.6 | L186I - N66D | -0.1972 | 1.7 | 239 | -0.116 | 1 |
| 0.6 | L186I - S167I | -1.7147 | 1.7 | 239 | -1.008 | 0.8516 |
| 0.6 | L186I - Triple | -2.8869 | 1.7 | 239 | -1.697 | 0.4379 |
| 0.6 | L186I - WT | -1.168 | 1.7 | 239 | -0.687 | 0.9593 |
| 0.6 | N66D - S167I | -1.5175 | 1.7 | 239 | -0.892 | 0.8997 |
| 0.6 | N66D - Triple | -2.6898 | 1.7 | 239 | -1.581 | 0.5111 |
| 0.6 | N66D - WT | -0.9709 | 1.7 | 239 | -0.571 | 0.9792 |
| 0.6 | S167I - Triple | -1.1722 | 1.7 | 239 | -0.689 | 0.9587 |
| 0.6 | S167I - WT | 0.5467 | 1.7 | 239 | 0.321 | 0.9977 |
| 0.6 | Triple - WT | 1.7189 | 1.7 | 239 | 1.01 | 0.8505 |

**Supplementary Table 4.** Difference in lytic activity between Lys1358 and the Triple variant (N66D, S167I, L186I) across enzyme concentrations in *Lactococcus lactis* 582. Estimated differences in lytic activity were obtained from estimated marginal means (EMMs)-based contrasts of a linear mixed-effects model at each enzyme concentration (μM). SE, standard error; df, degrees of freedom; t ratio, test statistic from the model contrasts; P value, significance of the comparison. Statistical significance is indicated as follows: *P < 0.05; **P < 0.01; ***P < 0.001.

| Concentration (μM) | Estimate | SE | df | t ratio | P value | Significance |
|---|---|---|---|---|---|---|
| 0.05 | -8.81824 | 1.460493 | 79 | -6.03785 | < 0.0001 | *** |
| 0.08 | -12.7289 | 1.460493 | 79 | -8.71551 | < 0.0001 | *** |
| 0.1 | -19.8376 | 1.460493 | 79 | -13.5828 | < 0.0001 | *** |
| 0.15 | -21.336 | 1.460493 | 79 | -14.6088 | < 0.0001 | *** |
| 0.2 | -23.8183 | 1.460493 | 79 | -16.3084 | < 0.0001 | *** |
| 0.3 | -20.4402 | 1.460493 | 79 | -13.9954 | < 0.0001 | *** |
| 0.45 | -14.3683 | 1.460493 | 79 | -9.83798 | < 0.0001 | *** |
| 0.6 | -10.8681 | 1.460493 | 79 | -7.4414 | < 0.0001 | *** |

**Supplementary Table 5.** Model-derived differences in ELISA binding activity between Lys1358 and variants across enzyme concentrations in *L. lactis* IL1403. Estimated differences in binding activity were obtained from estimated marginal means (EMMs)-based contrasts of a linear mixed-effects model fitted to absorbance values at each enzyme concentration (µM). Positive estimates indicate higher binding relative to Lys1358. Percent changes relative to Lys1358 were calculated from model-derived estimated marginal means (EMMs) as [(EMM_variant / EMM_Lys1358) × 100] − 100. The "Triple" construct corresponds to the combined N66D, S167I, and L186I substitutions. SE, standard error; df, degrees of freedom; t_ratio, test statistic from model contrasts; P_value, significance of the comparison. Statistical significance is indicated as follows: *P < 0.05; **P < 0.01; ***P < 0.001.

| Concentration (µM) | Construct | Estimate | SE | df | t ratio | % of Lys1358 | P value | Significance |
|---|---|---|---|---|---|---|---|---|
| 0.4 | L186I | 0.003 | 0.056 | 219 | 0.055 | 0.158 | 1 | |
| 0.4 | N66D | 0.721 | 0.056 | 219 | 12.876 | 36.951 | <0.001 | *** |
| 0.4 | S167I | 0.385 | 0.056 | 219 | 6.874 | 19.726 | <0.001 | *** |
| 0.4 | Triple | 0.751 | 0.056 | 219 | 13.421 | 38.515 | <0.001 | *** |
| 0.04 | L186I | 0.08 | 0.056 | 219 | 1.43 | 16.045 | 0.413 | |
| 0.04 | N66D | 2.165 | 0.056 | 219 | 38.682 | 434.18 | <0.001 | *** |
| 0.04 | S167I | 0.138 | 0.056 | 219 | 2.46 | 27.612 | 0.051 | |
| 0.04 | Triple | 1.699 | 0.056 | 219 | 30.352 | 340.682 | <0.001 | *** |
| 0.004 | L186I | 0.054 | 0.056 | 219 | 0.957 | 55.051 | 0.71 | |
| 0.004 | N66D | 1.024 | 0.056 | 219 | 18.29 | 1051.627 | <0.001 | *** |
| 0.004 | S167I | 0.27 | 0.056 | 219 | 4.832 | 277.825 | <0.001 | *** |
| 0.004 | Triple | 2.494 | 0.056 | 219 | 44.558 | 2561.901 | <0.001 | *** |
| 0.0004 | L186I | 0.019 | 0.056 | 219 | 0.344 | 28.589 | 0.972 | |
| 0.0004 | N66D | 0.053 | 0.056 | 219 | 0.952 | 79.084 | 0.714 | |
| 0.0004 | S167I | 0.043 | 0.056 | 219 | 0.762 | 63.366 | 0.821 | |
| 0.0004 | Triple | 0.545 | 0.056 | 219 | 9.746 | 810.025 | <0.001 | *** |

**Supplementary Table 6.** Model-derived differences in ELISA binding activity between Lys1358 and variants across enzyme concentrations in *L. lactis* 582. Estimated differences in binding activity were obtained from estimated marginal means (EMMs)-based contrasts of a linear mixed-effects model fitted to absorbance values at each enzyme concentration (µM). Positive estimates indicate higher binding relative to Lys1358. Percent changes relative to Lys1358 were calculated from model-derived estimated marginal means (EMMs) as [(EMM_variant / EMM_Lys1358) × 100] − 100. The "Triple" construct corresponds to the combined N66D, S167I, and L186I substitutions. SE, standard error; df, degrees of freedom; t_ratio, test statistic from model contrasts; P_value, significance of the comparison. Statistical significance is indicated as follows: *P < 0.05; **P < 0.01; ***P < 0.001.

| Concentration (µM) | Construct | Estimate | SE | df | t ratio | % of Lys1358 | P value | Significance |
|---|---|---|---|---|---|---|---|---|
| 0.4 | L186I | -0.476 | 0.091 | 219 | -5.211 | -20.284 | <0.001 | *** |
| 0.4 | N66D | 0.029 | 0.091 | 219 | 0.316 | 1.23 | 0.977 | |
| 0.4 | S167I | 0.072 | 0.091 | 219 | 0.784 | 3.053 | 0.81 | |
| 0.4 | Triple | 0.402 | 0.091 | 219 | 4.4 | 17.125 | <0.001 | *** |
| 0.04 | L186I | -0.497 | 0.091 | 219 | -5.447 | -31.226 | <0.001 | *** |
| 0.04 | N66D | 0.514 | 0.091 | 219 | 5.634 | 32.299 | <0.001 | *** |
| 0.04 | S167I | -0.495 | 0.091 | 219 | -5.419 | -31.069 | <0.001 | *** |
| 0.04 | Triple | 0.412 | 0.091 | 219 | 4.515 | 25.887 | <0.001 | *** |
| 0.004 | L186I | -0.466 | 0.091 | 219 | -5.103 | -45.89 | <0.001 | *** |
| 0.004 | N66D | 0.145 | 0.091 | 219 | 1.589 | 14.287 | 0.324 | |
| 0.004 | S167I | -0.21 | 0.091 | 219 | -2.301 | -20.691 | 0.077 | |
| 0.004 | Triple | 0.721 | 0.091 | 219 | 7.898 | 71.032 | <0.001 | *** |
| 0.0004 | L186I | -0.146 | 0.091 | 219 | -1.599 | -48.558 | 0.319 | |
| 0.0004 | N66D | -0.031 | 0.091 | 219 | -0.336 | -10.205 | 0.973 | |
| 0.0004 | S167I | -0.073 | 0.091 | 219 | -0.797 | -24.21 | 0.803 | |
| 0.0004 | Triple | 0.308 | 0.091 | 219 | 3.371 | 102.385 | 0.003 | ** |

**Supplementary Table 7.** X-ray data collection and refinement statistics; values in parentheses correspond to the highest-resolution shell.

| **Structure** | **Lys1358** | **Lys1358 (Leu186 alternate conf)** | **Lys1358_N66D** |
|---|---|---|---|
| **Space group** | **C2** | **C2** | **I2** |
| *a, b, c* (Å)<br>α, β, γ (°) | 123.7, 39.5, 56.1<br>90, 108.8, 90 | 123.8, 39.7, 56.1<br>90, 108.8, 90 | 55.7, 39.3, 118.4<br>90, 97.5, 90 |
| wavelength(Å) | 0.97931 | 0.97931 | 1.0332 |
| resolution (Å) | 58.6-1.33 (1.40-1.33) | 58.6-1.22 (1.29-1.22) | 47.7-1.52 (1.55-1.52) |
| observed *hkl* | 181145 (26511) | 233778 (23022) | 266465 (13182) |
| unique *hkl* | 55829 (8326) | 72281 (9125) | 39411 (1921) |
| redundancy | 3.2 (3.2) | 3.2 (2.5) | 6.8 (6.9) |
| completeness (%) | 94.5 (96.5) | 94.3 (82.3) | 99.9 (100) |
| $R_{meas}$ | 0.038 (0.626) | 0.030 (0.216) | 0.049 (1.077) |
| $CC_{1/2}$ | 0.999 (0.793) | 0.999 (0.979) | 1.000 (0.837) |
| I/(σI) | 15.9 (2.2) | 22.5 (5.2) | 19.2 (2.0) |
| Wilson B (Å$^2$) | 15.1 | 12.2 | 18.3 |
| $R_{work}$ (# *hkl*) | 0.170 (52956) | 0.161 (68721) | 0.173 (37416) |
| $R_{free}$ (# *hkl*) | 0.196 (2844) | 0.175 (3555) | 0.210 (1971) |

| | | | |
|---|---|---|---|
| B-factors ($Å^2$)<br>(# atoms)<br>Protein<br>Ligands<br>Water | 21.0 (1749)<br>29.5 (19)<br>33.9 (257) | 16.5 (1747)<br>28.9 (29)<br>28.3 (283) | 25.3 (1733)<br>47.1 (61)<br>37.7 (173) |
| Ramachandran | | | |
| favored (%) | 100 | 100 | 100 |
| Generally allowed (%) | 0 | 0 | 0 |
| outlier(%) | 0 | 0 | 0 |
| rmsd | | | |
| bonds (Å) | 0.0149 | 0.0205 | 0.0124 |
| angles (°) | 1.885 | 2.157 | 1.700 |
| PDB code | 10VD | 12LL | 10VE |

**Supplementary Table 8.** Bacterial strains, bacteriophages, plasmids, and nucleotides used in this study.

| **Bacterial strains** | **Characteristics** | **Origin** |
|---|---|---|
| ***E. coli*** | | |
| BL21(DE3) | F$^-$ *ompT hsdSB* ($r_B^-$ $m_B^-$) *dcm*$^+$*gal* (DE3) | Stratagene |
| BL21/pLys1358$^{28a}$ | *E. coli* BL21 (DE3) transformed with pLys1358$^{28a}$ | [1] |
| BL21/pLys1358 N66D$^{28a}$ | *E. coli* BL21 (DE3) transformed with pLys1358 N66D$^{28a}$ | This study |
| BL21/pLys1358 S167I$^{28a}$ | *E. coli* BL21 (DE3) transformed with pLys1358 S167I$^{28a}$ | This study |
| BL21/pLys1358 L186I$^{28a}$ | *E. coli* BL21 (DE3) transformed with pLys1358 L186I$^{28a}$ | This study |
| BL21/pLys1358 N66D, S167I, L186I$^{28a}$ | *E. coli* BL21 (DE3) transformed with pLys1358 N66D, S167I, L186I$^{28a}$ | This study |
| BL21/pLys1358 C29A$^{28a}$ | *E. coli* BL21 (DE3) transformed with pLys1358 C29A$^{28a}$ | This study |
| BL21/pLys1358 C29A, N66D$^{28a}$ | *E. coli* BL21 (DE3) transformed with pLys1358 C29A, N66D$^{28a}$ | This study |
| BL21/pLys1358 C29A, S167I$^{28a}$ | *E. coli* BL21 (DE3) transformed with pLys1358 C29A, S167I$^{28a}$ | This study |
| BL21/pLys1358 C29A, L186I$^{28a}$ | *E. coli* BL21 (DE3) transformed with pLys1358 C29A, L186I$^{28a}$ | This study |
| BL21/pLys1358 C29A, N66D, S167I, L186I $^{28a}$ | *E. coli* BL21 (DE3) transformed with pLys1358 C29A, N66D, S167I, L186I $^{28a}$ | This study |
| ***L. lactis*** | | |
| IL1403 | Laboratory strain, host of phage P008 | [2] |
| 582 | Host of phage 1358 | [3] |

| **Phages** | **Characteristics** | **Origin** |
|---|---|---|
| P008 | Phage infecting *L. lactis* IL1403 | [4] |
| 1358 | Phage infecting *L. lactis* 582 | [5] |
| P008::Lys1358 | Chimeric lactococcal phage derived from P008 in which the native endolysin gene (*lysP008*) is replaced by *lys1358*, encoding the endolysin of phage 1358. | [1] |

| Plasmids | Characteristics[a] | Origin |
|---|---|---|
| pET-28a | Expression vector; Kan$^r$ | Novagen |
| pLys1358$^{28a}$ | pET-28a carrying *Lys1358* | This study |
| pLys1358 N66D$^{28a}$ | pET-28a carrying *Lys1358* N66D | This study |

| | | |
|---|---|---|
| pLys1358 S167I[28a] | pET-28a carrying *Lys1358* S167I | This study |
| pLys1358 L186I[28a] | pET-28a carrying *Lys1358* L186I | This study |
| pLys1358 N66D, S167I, L186I[28a] | pET-28a carrying *Lys1358* N66D, S167I, L186I | This study |
| pLys1358 C29A[28a] | pET-28a carrying *Lys1358* C29A | This study |
| pLys1358 C29A, N66D[28a] | pET-28a carrying *Lys1358* C29A, N66D | This study |
| pLys1358 C29A, S167I[28a] | pET-28a carrying *Lys1358* C29A, S167I | This study |
| pLys1358 C29A, L186I[28a] | pET-28a carrying *Lys1358* C29A, L186I | This study |
| pLys1358 C29A, N66D, S167I, L186I [28a] | pET-28a carrying *Lys1358* C29A, N66D, S167I, L186 | This study |

| Oligonucleotides | Sequence 5’ → 3’ | Origin |
|---|---|---|
| Lys1358_N66D_Fw | ACGCATCAAGGACAGCCGGGGGT | This study |
| Lys1358_N66D_Rv | TGCCAGCCCGCCGGAAG | This study |
| Lys1358_S167I_Fw | CAAAGCAAACATTGCCGTCAACATCCGTC | This study |
| Lys1358_S167I _Rv | AACGTCCCCCGCTGA | This study |
| Lys1358_L186I_Fw | AGCCGGGGTTATTAAAGCAGGCC | This study |
| Lys1358_L186I_Rv | ACGGTACCTGTTTTTGTGTTC | This study |
| Lys1358_C29A_Fw | CGGTGCACAGGCGATGGACCTCG | This study |
| Lys1358_C29A _Rv | TACATGCCGTCAAAATC | This study |
| T7 | TAATACGACTCACTATAGGG | Novagen |
| T7 terminator | GCTAGTTATTGCTCAGCGG | Novagen |

[a] Abbreviations: $Cam^r$, chloramphenicol resistance; $Kan^r$, kanamycin resistance [b]

**Supplementary Table 1 references:**
