## Supplementary Figures for "Predictable adaptive evolution of a phage endolysin through substrate recognition optimization"

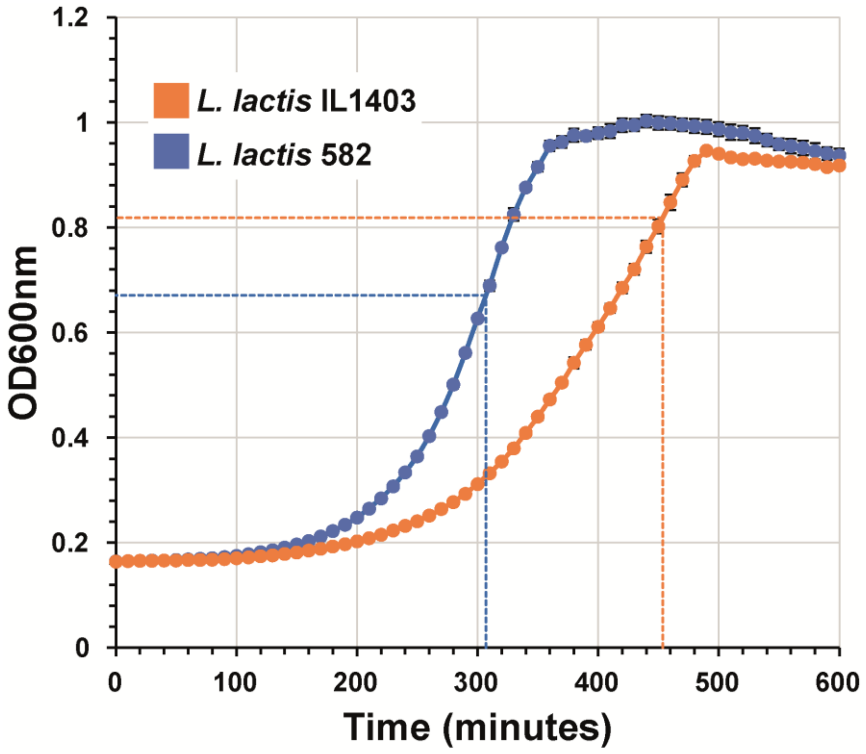


**Supplementary Fig. 1 |** Growth curves of *L. lactis* IL1403 and 582. Cells were cultured in 96-well plates, and $OD_{600}$ was recorded every 10 min over 960 min. The time to maximum growth rate was 454.3 ± 5.8 min for strain 582 and 307.7 ± 5.8 min for IL1403. Experiments were performed in triplicate.

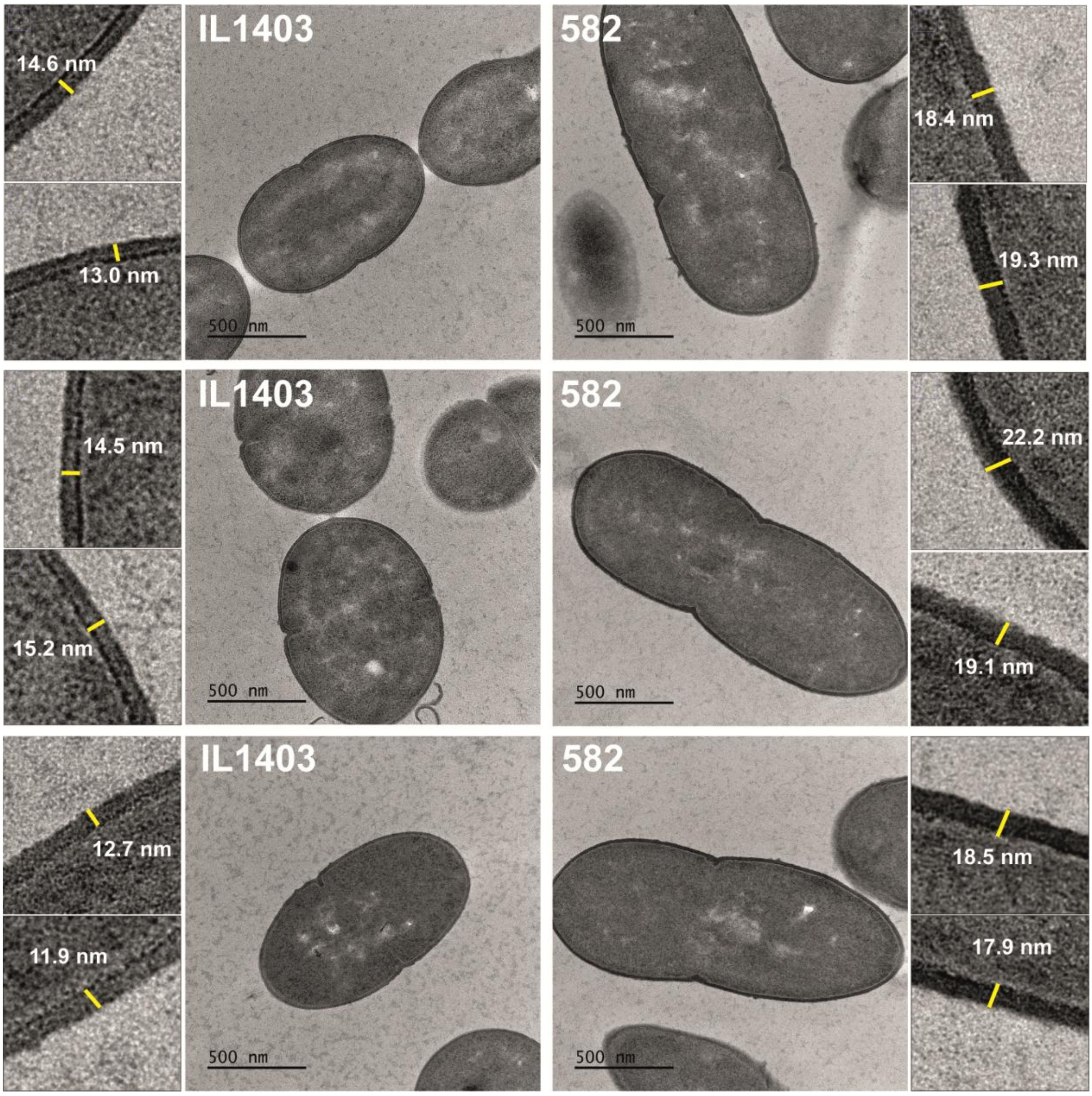


**Supplementary Fig. 2 |** Cell wall thickness of *L. lactis* strains IL1403 (13.6 ± 1.3 nm) and 582 (19.2 ± 1.5 nm), determined by transmission electron microscopy (TEM).

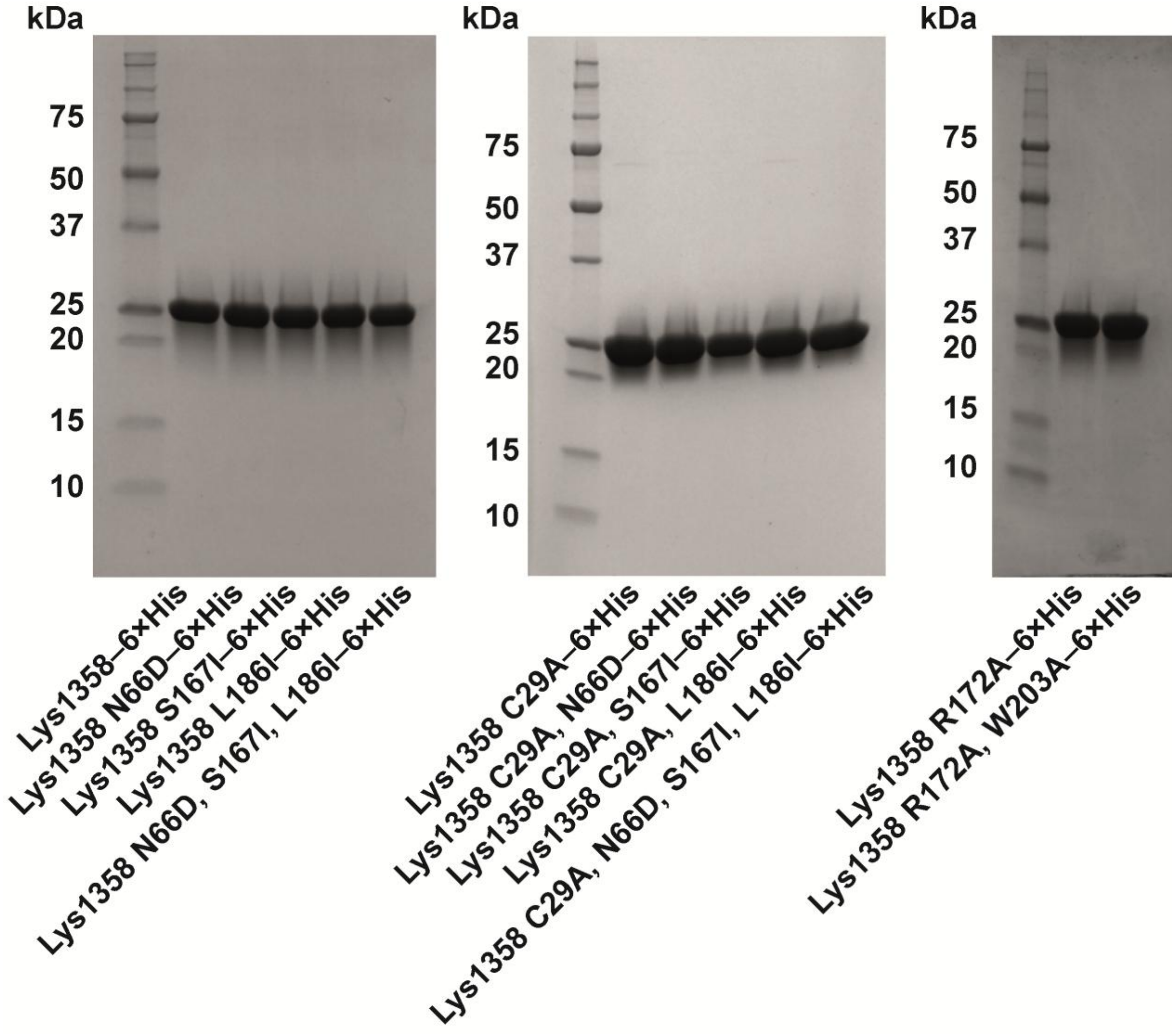


**Supplementary Fig. 3 |** SDS–PAGE gel showing the purified endolysin constructs used in the study. Proteins were separated on NuPAGE 4–12% Bis–Tris gels and visualized by Coomassie Blue staining. Molecular weight markers are shown on the left of each gel.

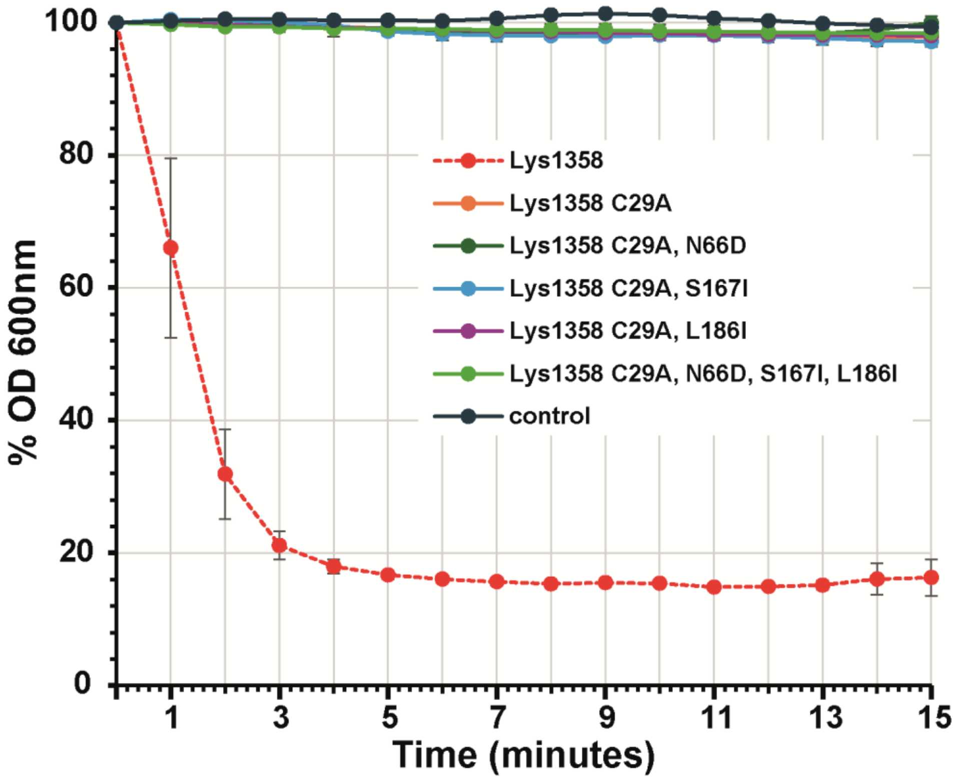


**Supplementary Fig. 4 |** Lytic activity of Lys1358 and C29A variants at 4 μM. Activity was assessed by monitoring the decrease in turbidity of *L. lactis* IL1403 cells in exponential phase at 1-min intervals over 15 min. Experiments were performed in triplicate.

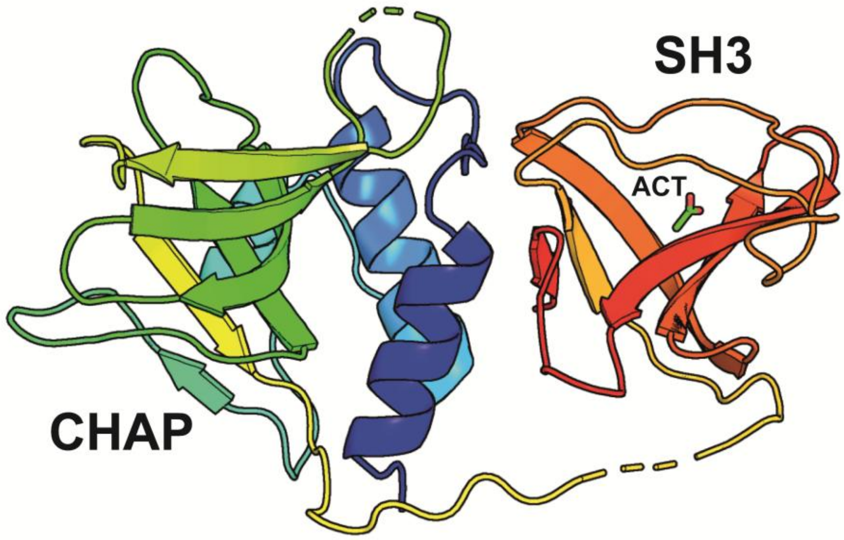


**Supplementary Fig. 5 |** Crystal structure of Lys1358 (PDB: 10VD). The structure was determined in space group C2 with one molecule in the asymmetric unit. All residues were modelled except for regions 109–115 and 147–152. The protein is shown in a rainbow representation, with the N-terminus in blue and the C-terminus in red. The CHAP catalytic domain (residues 1–139) and the SH3 cell wall-binding domain (residues 155–280) are indicated. An acetate molecule (ACT) bound to the CBD domain is also shown. domain (residues 155–280) are indicated. An acetate molecule (ACT) bound to the CBD domain is also shown.

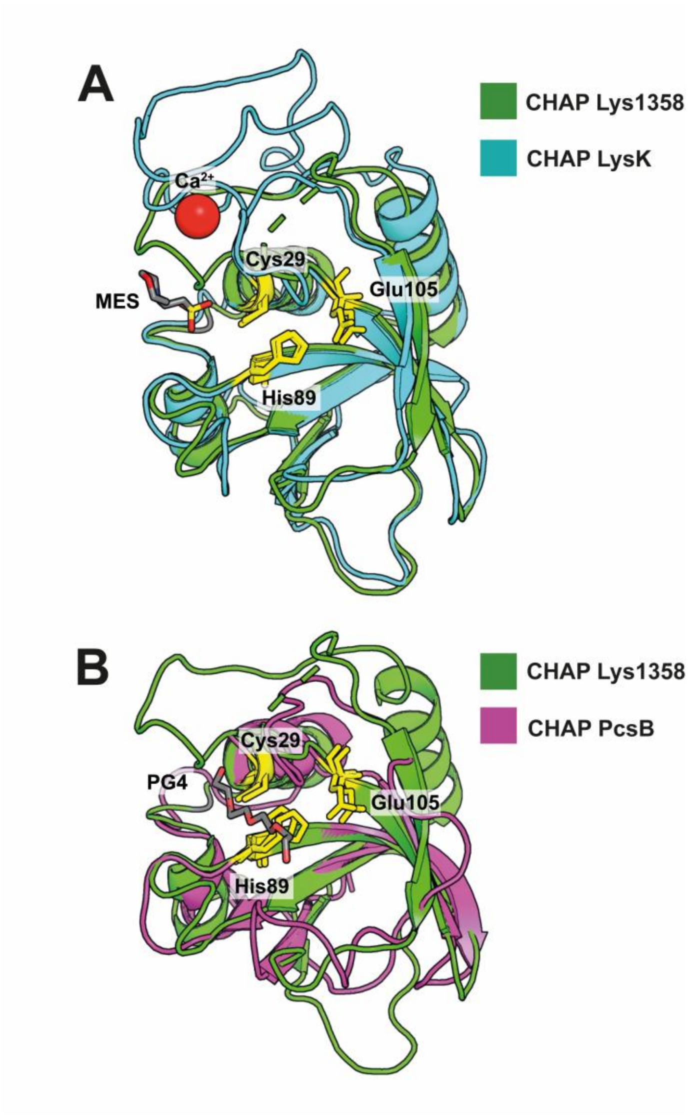


**Supplementary Fig. 6 |** Superposition of the Lys1358 CHAP domain (residues 1–139, green) with structural homologues. **(a)** CHAP domain of the LysK endolysin (residues 1–165, PDB ID: 4CSH, light magenta). **(b)** CHAP domain of the ParB autolysin (residues 291–391, PDB ID: 4CGK, light purple). The catalytic residues of Lys1358 (Cys29, His89, Glu105) are shown as yellow sticks, with the corresponding conserved residues in CHAPk (Cys54, His117, Glu134) and ParB (Cys292, His343, Glu360) indicated. Co-crystallized ligands—2-(N-morpholino)ethanesulfonic acid (MES) for CHAPk and tetraethylene glycol (PG4) for ParB—highlight the catalytic site and are shown in red.

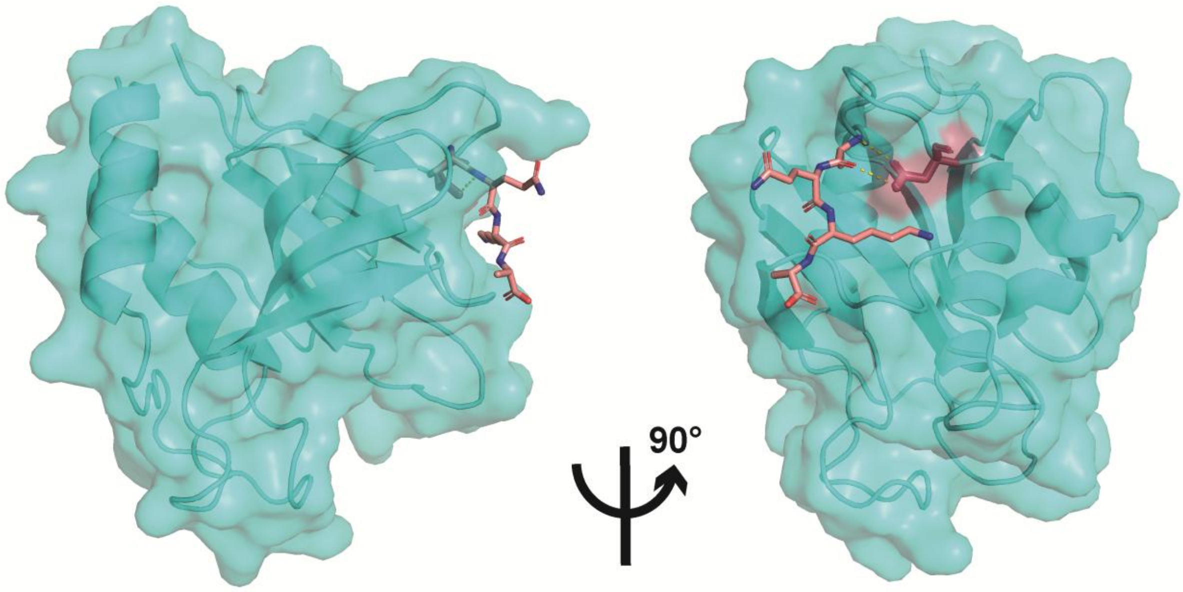


**Supplementary Fig. 7 |** AlphaFold prediction of the Lys1358 CHAP domain bound to the AEQA peptide. The N66 residue is highlighted in red. Yellow lines indicate the closest approach between N66 and the alanine residue of the AEQA peptide, illustrating their spatial proximity.

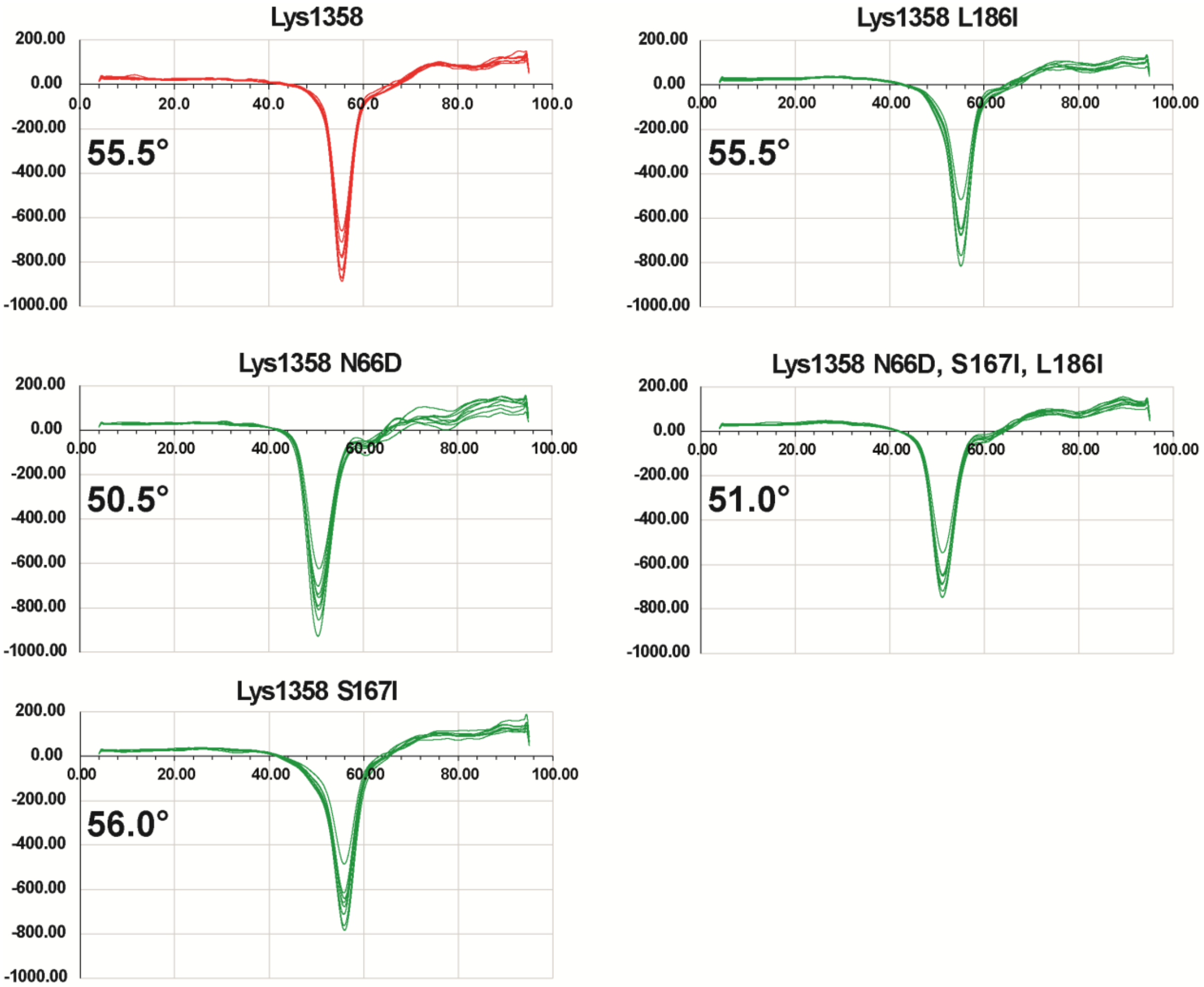


**Supplementary Fig. 8 |** Melting temperatures of Lys1358 and its variants. Thermal shift assays were performed using SYPRO® Orange, and the melting temperature ($T_m$) was determined from the minimum of the first derivative of fluorescence intensity with respect to temperature (0.5 °C increments).

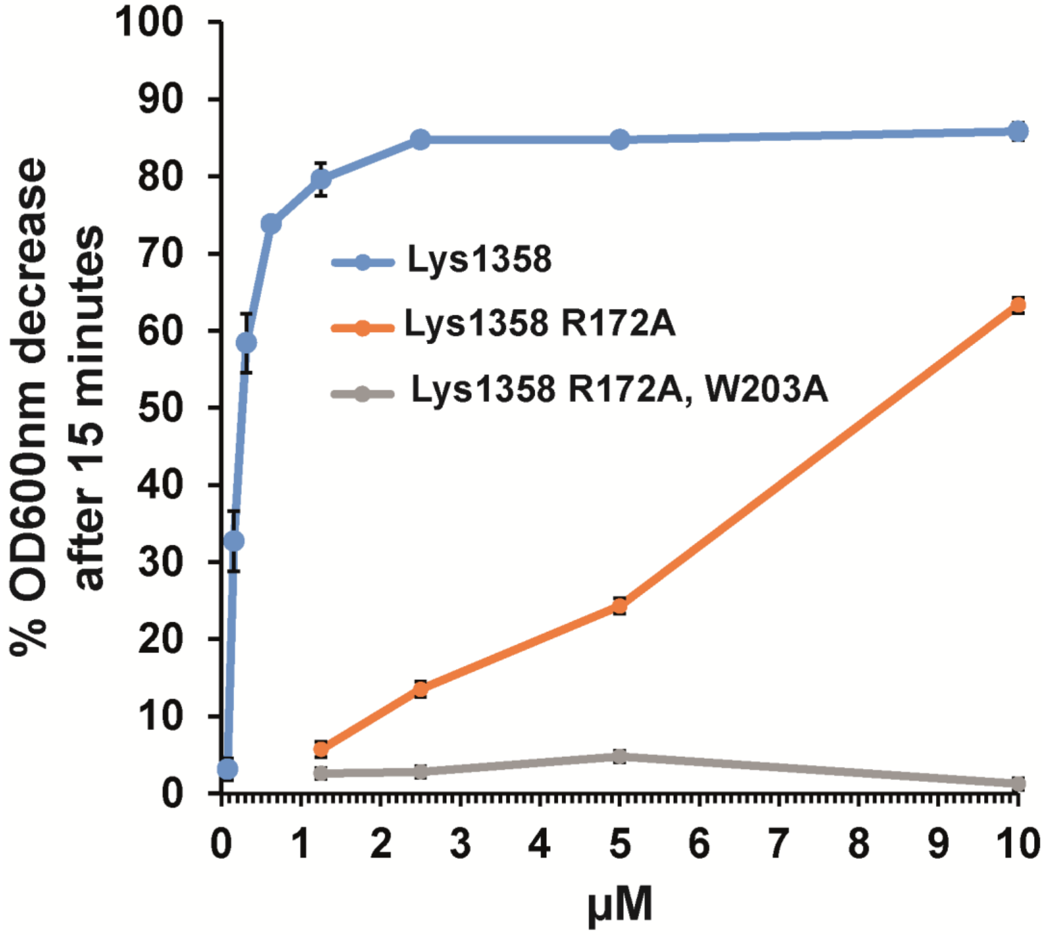


**Supplementary Fig. 9 |** Lytic activity of Lys1358, the R172A mutant, and the R172A/W203A double mutant. Activity was assessed at multiple concentrations by monitoring the decrease in turbidity of *L. lactis* IL1403 cells in exponential phase. Turbidity at 600 nm was measured after 15 min of incubation and expressed as percent reduction.

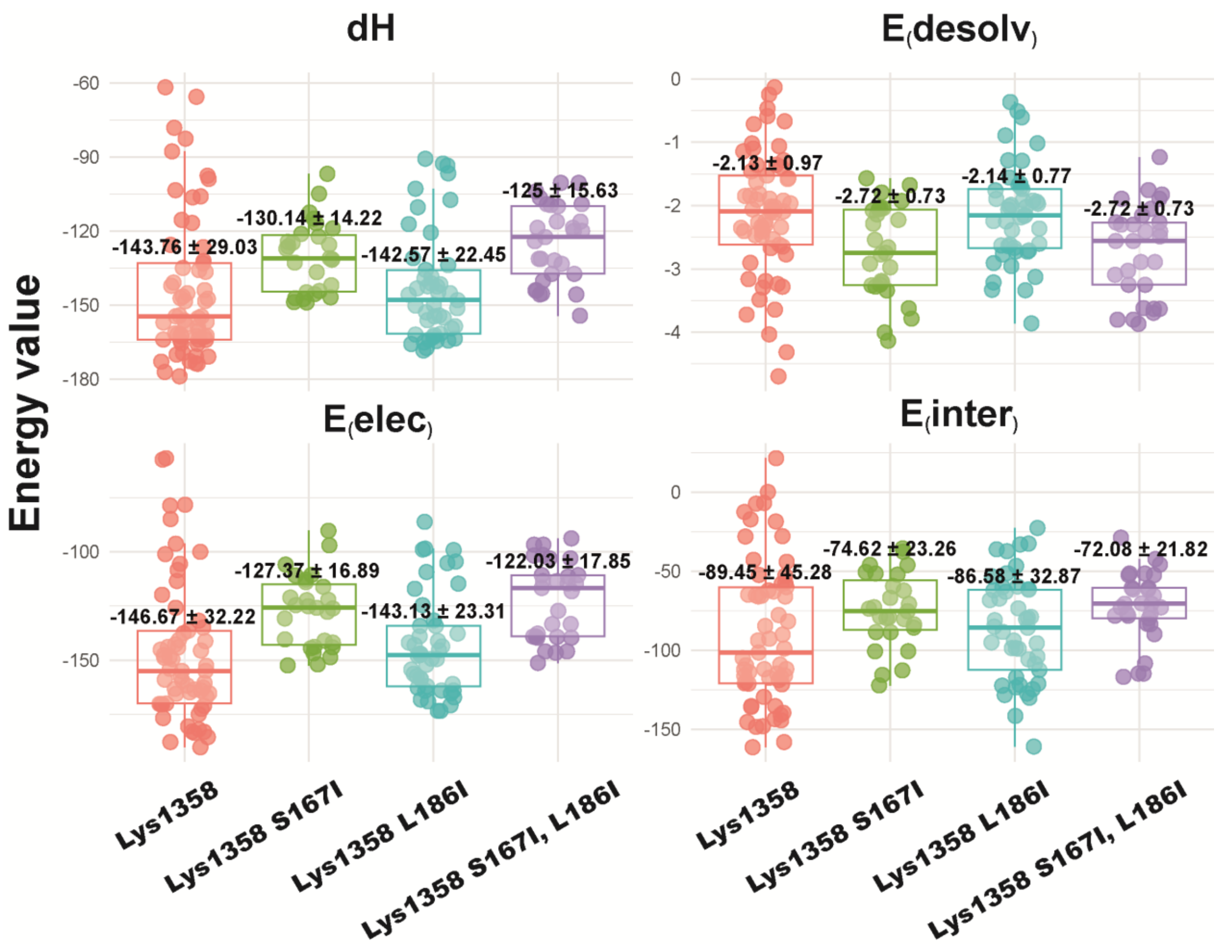


**Supplementary Fig. 10 |** HADDOCK-derived energy values for the SH3 domain of Lys1358 (wild-type) and its variants (S167I, L186I, and the double variant S167I/L186I) in complex with the AKQA peptide. The peptide was taken from the crystal structure of the lysostaphin SH3 domain (PDB ID: 6RK4), where it was originally co-crystallized. Box plots show the distribution of energy terms: dH (HADDOCK score combining van der Waals, electrostatics, and desolvation contributions), E(desolv) (desolvation), E(elec) (electrostatics), and E(inter) (intermolecular energy). Energies were calculated using HADDOCK 2.4 with default parameters and are expressed in arbitrary HADDOCK units. See Methods for details.

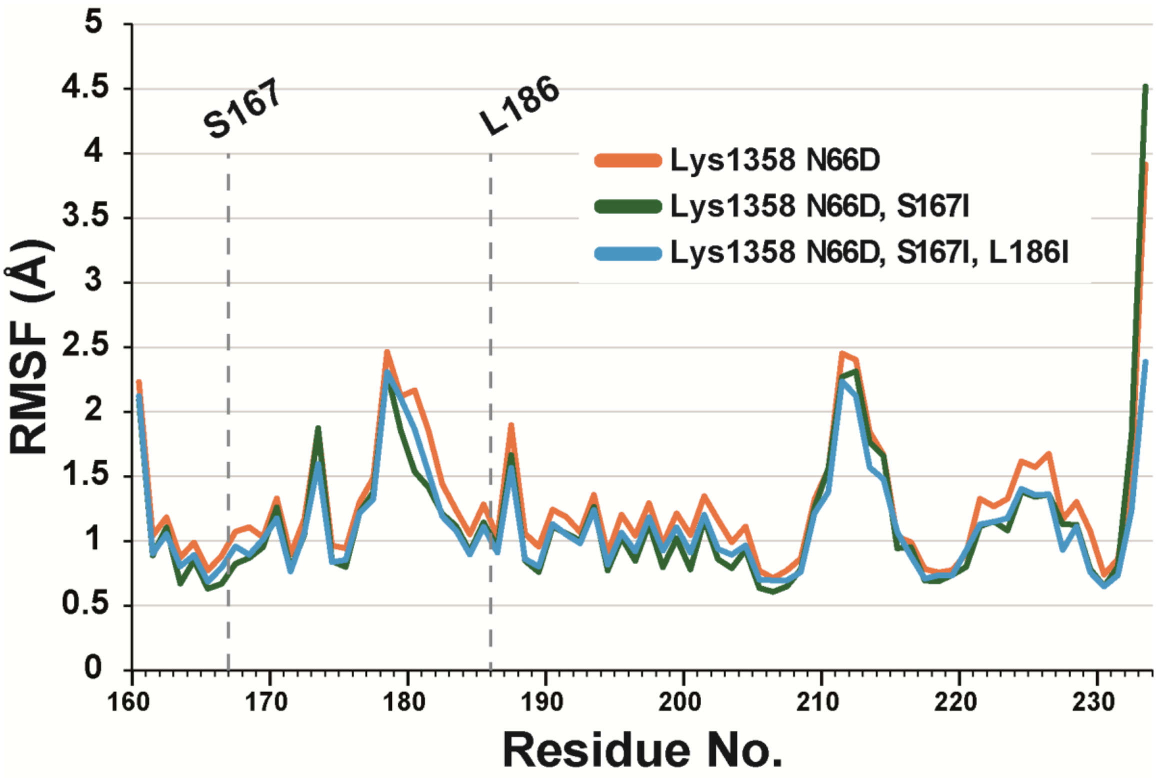


**Supplementary Fig. 11 |** Molecular dynamics simulations of Lys1358 variants: N66D (orange), N66D/S167I (green), and N66D/S167I/L186I (blue). Only residues 160–233, corresponding to the SH3 domain of Lys1358, are shown to highlight the region containing the mutations. Simulations were performed as described in Methods.

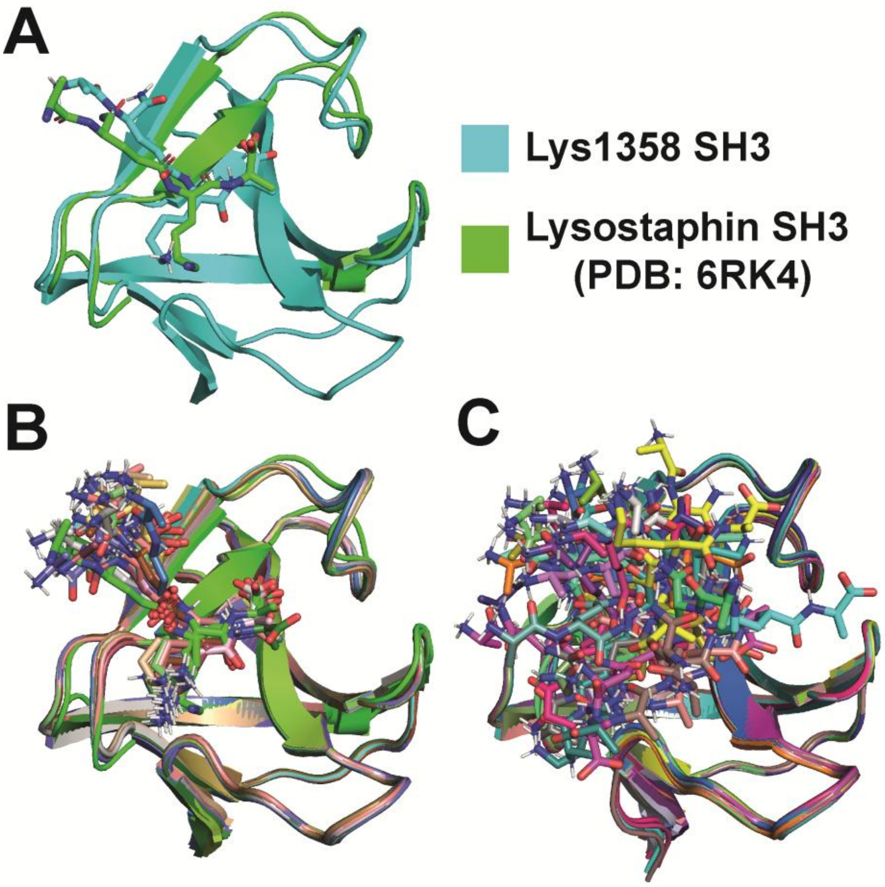


**Supplementary Fig. 12 |** Docking simulations of the Lys1358 SH3 domain with the AQKA stem peptide. The peptide sequence was derived from the crystal structure of the lysostaphin SH3 domain (PDB ID: 6RK4), where it was originally co-crystallized. Docking was performed using HADDOCK 2.4 with default parameters. The first 50 models (of 200) illustrate predicted interactions between the Lys1358 SH3 domain and the AQKA stem peptide. **(a)** Example of a correctly oriented complex showing a peptide conformation consistent with that observed in lysostaphin. **(b)** Representative correctly oriented complexes among the first 50 models. **(c)** Representative misoriented complexes among the first 50 models; these were excluded from subsequent energy analyses (Supplementary Fig. 10).
